# Selective control of working memory in prefrontal, parietal, and visual cortex

**DOI:** 10.1101/2020.04.07.030718

**Authors:** Matthew F. Panichello, Timothy J. Buschman

## Abstract

Cognitive control guides behavior by controlling what, where, and how information is represented in the brain. Previous work has shown parietal and prefrontal cortex direct attention, which controls the representation of external sensory stimuli^1,2^. However, the neural mechanisms controlling the selection of representations held ‘in mind’, in working memory, are unknown. To address this, we trained two monkeys to switch between two tasks, requiring them to either select an item from a set of items held in working memory or attend to one stimulus from a set of visual stimuli. Simultaneous neural recordings in prefrontal, parietal, and visual cortex found prefrontal cortex played a primary role in selecting an item from working memory, representing selection before parietal and visual cortex. Surprisingly, a common population representation in prefrontal cortex encoded selection of an item in working memory and attention to an external stimulus, suggesting prefrontal cortex may act as a domain-general controller. Selection acted on memory representations in two ways. First, selection improved the accuracy of memory reports by enhancing the selected item’s representation in prefrontal and parietal cortex. Second, selection transformed memory representations in a task-dependent manner. Before selection, when both items were relevant to the task, the identity of each item was represented in an independent subspace of neural activity. After selection, the representation of only the selected item was transformed into a new subspace that was used to guide the animal’s behavioral report. Together, our results provide insight into how prefrontal cortex controls working memory representations, selectively enhancing and transforming them to support behavior.

## Main Text

Working memory maintains information relevant to the current task (e.g. the list of specials at a restaurant). Items held in working memory are thought to be represented in a distributed network of brain regions, including prefrontal cortex, parietal cortex, and sensory cortex^3^. A control mechanism can then select a specific item from working memory and use it to guide behavior^4–7^ (e.g. selecting a special to order for dinner). This process is similar to attention, which selectively enhances task-relevant sensory inputs^1,2^. Previous functional imaging work has shown prefrontal and parietal cortex are active when an item is selected from working memory^8–10^. However, because it has never been studied at the level of single neurons, the neural mechanisms of selection remain unknown.

To address this, we simultaneously recorded from the prefrontal, parietal, and visual cortices of two monkeys (*Macaca mulatta*) as they selected one of two items held in working memory. On each trial of the experiment, the animals remembered the color of two squares (Fig. 1A, an ‘upper’ and ‘lower’ stimulus). After a memory delay, the animals received a cue that indicated whether they should report the color of the ‘upper’ or ‘lower’ square (now held in working memory). This cue was followed by a second memory delay, after which the animals reported the color of the cued square by saccading to the matching color on a color wheel (note, the wheel was randomly rotated on each trial to prevent motor planning). Therefore, to perform the task, the animals had to hold two colors in working memory, select the color of the cued square, and then use it to guide their response to the color wheel.

**Figure 1.**
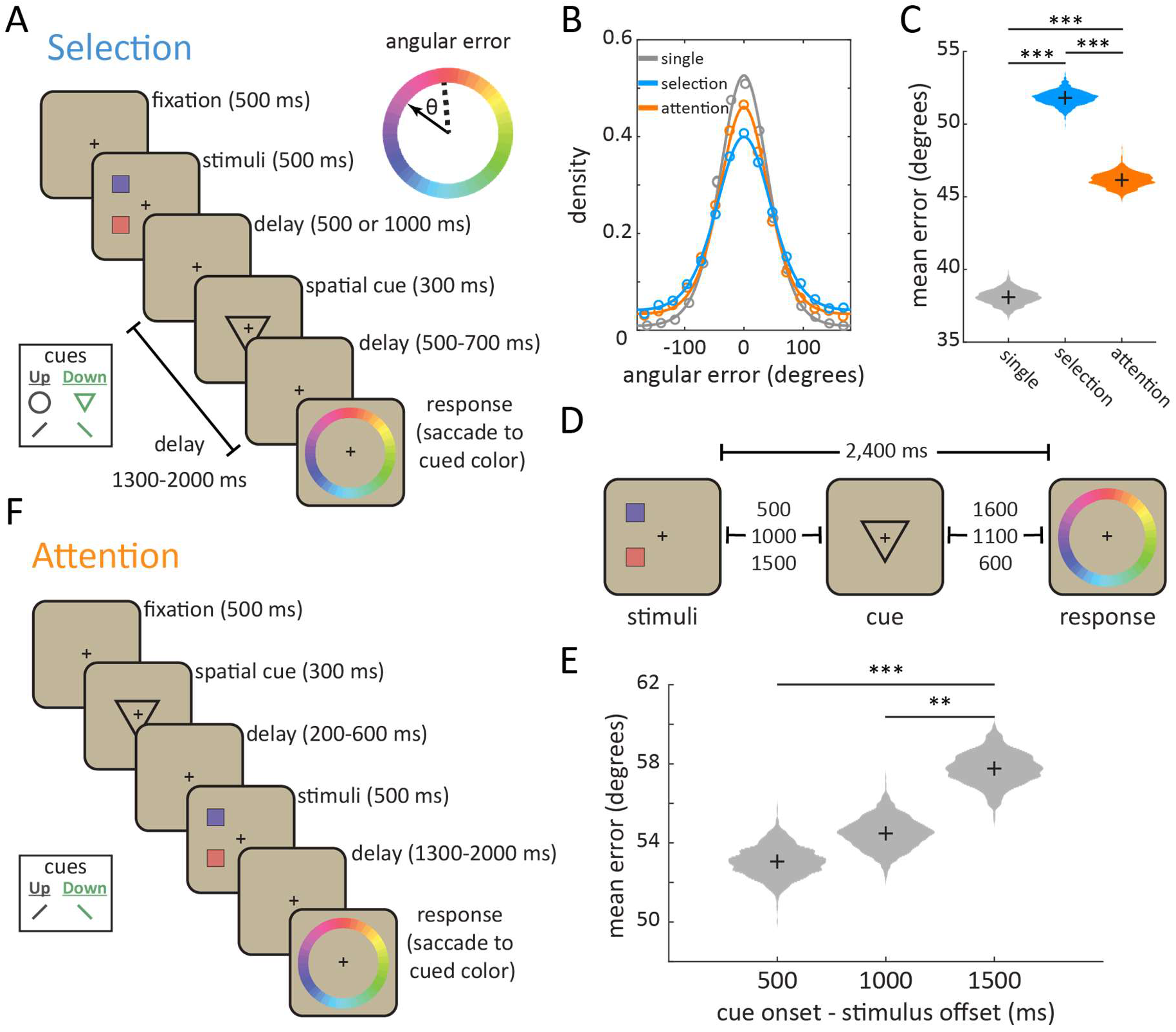
Monkeys use selection and attention to control the contents of working memory. **(A)** Animals were trained to perform two variants of a delayed estimation paradigm. On each trial, two colored squares were presented (one ‘upper’ and one ‘lower’ stimulus). Lower inset shows symbolic cues presented at fixation to indicate whether the upper or lower stimulus should be reported at the end of the trial in order to receive reward. In the ‘selection’ condition, the cue appeared during a memory delay after stimulus offset, requiring the animal to select an item from working memory. Animals received a graded juice reward for making an eye movement to the portion of a color wheel matching the color of the cued stimulus. Color wheel inset shows error was calculated as the angular deviation between the presented color (dashed) and the reported color (solid). **(B)** Distribution of angular error (circles) with best-fitting mixture models (lines, see Methods) for single item trials (gray), selection trials (blue), and attention trials (orange). **(C)** Bootstrapped distribution of mean angular error in the selection, attention, and single-stimulus conditions (colors as in B). **(D)** In a separate behavioral experiment, we fixed the total memory delay of the selection condition and systematically varied the length of the delay between stimuli offset and cue onset. **(E)** Increasing the time before selection increased error. Distribution shows mean angular error as a function of the delay between stimulus and selection cue (bootstrapped). Bars and asterisks reflect paired randomization tests. **(F)** Animals also performed an ‘attention’ condition, interleaved with selection trials (in blocks). In this condition, the cue appeared before stimulus onset. Inset: the two symbolic cues used in the attention condition. * p < 0.05, ** p < 0.01, *** p < 0.001.

Overall, both monkeys performed the task well; the mean angular error between the presented and reported color was ~50° (43.8° and 58.5° for monkey 1 and 2, respectively, Fig. S1). As expected^11–13^, accuracy depended on the number of items in memory (i.e. the ‘memory load’) – angular error was greater when two colored squares were presented, compared to a separate set of trials in which only one square was presented (Fig. 1B-C and S2A; 38.1° mean absolute error for 1 item, 51.8° for 2 items, p < 0.001, randomization test). The increased error with two items in memory is thought to be due to interference between the items held in working memory^14–17^Λ Selecting an item in working memory is thought to reduce this interference^18,19^. Consistent with this, reports were more accurate when selection occurred earlier in the trial (Fig. 1D-E, 53.1°, 54.4°, and 57.8° for 0.5, 1, and 1.5 s post-cue, respectively; linear regression, β=4.67 ± 1.08 SEM, p< 0.001, bootstrap). Behavioral modeling^12,13^ showed earlier cues improved the precision of memory reports (Fig. S2B, β=3.95 ± 1.88 SEM, p = 0.012, bootstrap) but did not significantly change the probability of forgetting (i.e. random responses; Fig. S2B, β=0.03 ± 0.03 STE, p = 0.126, bootstrap).

In addition to the selection condition, animals also performed an ‘attention’ condition. On attention trials, the cue was presented before the colored squares, allowing the animal to attend to the location of the to-be-reported stimulus (Fig. 1F). Memory reports were more accurate in the attention condition than in the selection condition (Fig. 1B-C; 46.1° vs. 51.8°, p < 0.001, randomization test). Behavioral modeling showed this was due to an increase in the precision of memory reports and a reduction in forgetting (i.e. fewer random reports; Fig. S2B). In addition, the effect of memory load was reduced; when the number of stimuli increased from 1 to 2 the mean absolute error increased by 9.01° on attention trials compared to 13.7° degrees on selection trials (p < 0.001, bootstrap). This is consistent with attention reducing interference between stimuli^1,20^ and modulating what enters working memory^21^.

To understand the neural mechanisms of selection, we simultaneously recorded from four regions known to be involved in working memory (Fig. 2A) – lateral prefrontal cortex (LPFC; 682 neurons), frontal eye fields (FEF; 187 neurons), parietal cortex (7a/b; 331 neurons), and intermediate visual area V4 (341 neurons). All four regions carried information about which item was selected from working memory (i.e. the upper or lower item). Figure 2B shows an example LPFC neuron that preferentially responds to selecting the upper item (see Fig. S3A-B for selectivity across the entire population). To quantify information about selection, we trained a logistic regression classifier to decode the location of selection from the firing rates of populations of neurons recorded in each region (Fig. 2C; pseudopopulations were constructed across all recording days, see methods for details). As seen in Figure 2D, the classifier could decode the location of selection in all four regions (blue lines). However, significant information about selection emerged first in LPFC and then later in posterior regions (Fig. 2E, 175 ms post-cue in LPFC, 245 ms in FEF, 285 ms in parietal, and 335 ms in V4). LPFC was significantly earlier than parietal cortex and V4 (p = 0.005 and p = 0.048, randomization test; but statistically indistinguishable from FEF, p=0.371). This suggests that signals necessary for controlling selection emerge first in prefrontal cortex (similar to prefrontal control of attention^22–24^).

**Figure 2.**
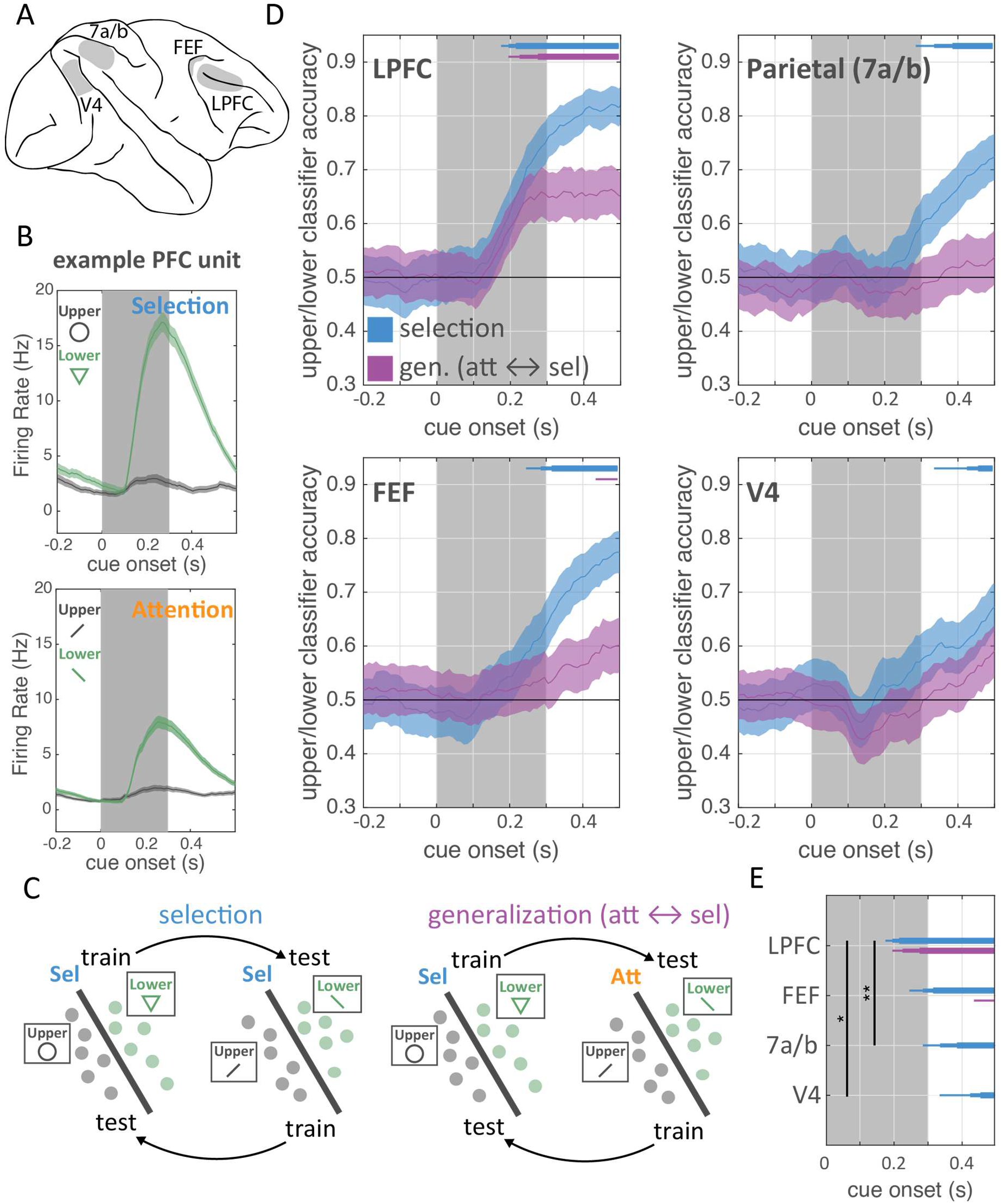
Selection is directed from prefrontal cortex and shares a population code with attention. **(A)** Neural activity was recorded simultaneously from lateral PFC (LPFC), the frontal eye fields (FEF), parietal cortex (area 7a/b), and visual area V4. **(B)** Firing rate of an example PFC neuron around cue onset when the upper (gray) or lower (green) stimulus was cued in the selection and attention conditions. Shaded regions are standard error of the mean. Inset shows different cues used for selection and attention. **(C)** Training and testing regime for classifiers designed to quantify information about the location of selection and attention based on population firing rates. Classification accuracy was measured on held-out data both within selection (left) and across selection and attention (right). Classifiers were trained and tested on different cue sets and performance was averaged across these splits. **(D)** Timecourse of information about the location of selection and attention for each brain region (labeled in upper left). Lines show mean classification accuracy around cue onset for the classifier trained and tested within the selection condition (blue) and across selection and attention (purple). Error bars are standard error of the mean. Bars along top indicate above-chance classification: p < 0.05, 0.01, and 0.001 for thin, medium, and thick lines, respectively. **(E)** Timepoints of significant classification for each ROI, as in (D). * p < 0.05, ** p < 0.01, *** p < 0.001.

There is a functional homology between selection and attention^4^. They both control neural representations – selection controls ‘internal’ working memory representations, while attention controls ‘external’ sensory representations. In both cases, control mitigates interference between representations (Fig. 1C-E,^1,14,25^). Motivated by their functional homology, we investigated whether there was a shared population code controlling selection and attention. Previous work in humans has shown both attention and selection activate prefrontal and parietal cortex^8–10,26^. However, it is not known if the neural mechanisms controlling selection and attention are the same. To test this, we first examined the responses of single neurons to the ‘upper’ and ‘lower’ cues on selection and attention trials. Neurons that encoded the location of selection responded similarly during attention in LPFC (Fig. S3C; r(586) = 0.09, p = 0.036). In contrast, sensitivity for selection and attention were uncorrelated in FEF, V4, and parietal cortex (Fig. S3C; FEF: r(169) = 0.04, p = 0.617; V4: r(318) = −0.04, p = 0.513; parietal: r(301) = 0.03, p = 0.612).

Furthermore, classifiers trained to decode the location of selection generalized to decode the location of attention (and vice versa; see Fig. 2C and Methods for details). Consistent with a common mechanism in LPFC, generalization performance was significantly above chance and followed the timecourse of the selection classifier (Fig. 2D, purple lines). In contrast, generalization was weaker in FEF and trended towards being delayed relative to LPFC (p = 0.12, randomization test), and failed to reach significance in parietal cortex and V4 (Fig. 2D-E; note, poor generalization was not due to an inability to decode the location of attention from these regions, Fig. S4). Together, these results suggest a common neural mechanism in LPFC controls attention to sensory inputs and selection of items in working memory.

Next, we explored how selection impacts the neural representations of items in working memory. As noted above, selection improves working memory accuracy^5,27–29^ (Fig. 1E). To understand the neural mechanisms of this improvement, we first measured the strength of color information in LPFC, FEF, parietal cortex (7a/b), and V4. Single neurons in all four regions showed strong color selectivity (see Fig. 3A for an example LPFC neuron). Selectivity was quantified by measuring the circular entropy of each neuron’s firing rate in response to colors across the color wheel (high entropy reflects high color information, see methods for details). A significant proportion of neurons in each region encoded color information about either the upper or lower stimulus during the trial (LPFC: N = 387/607 cells; FEF: 114/178; parietal: 181/307; V4: 245/323; all p < 0.001 by binomial test, corrected for multiple comparisons across time and stimulus location). Across the population, all four regions carried strong information about the color of the stimulus upon its presentation (Fig. 3B, left panels). Color information was then maintained across this distributed network during the first memory delay (Fig. 3B; middle panels, before selection). Interestingly, there was less information about color on attention trials compared to selection trials (Fig. S5–S6). This could reflect a task-specific difference in how memories are stored: recent theoretical work^30^ suggests a more active representation is needed when manipulating memories (such as in the selection condition).

**Figure 3.**
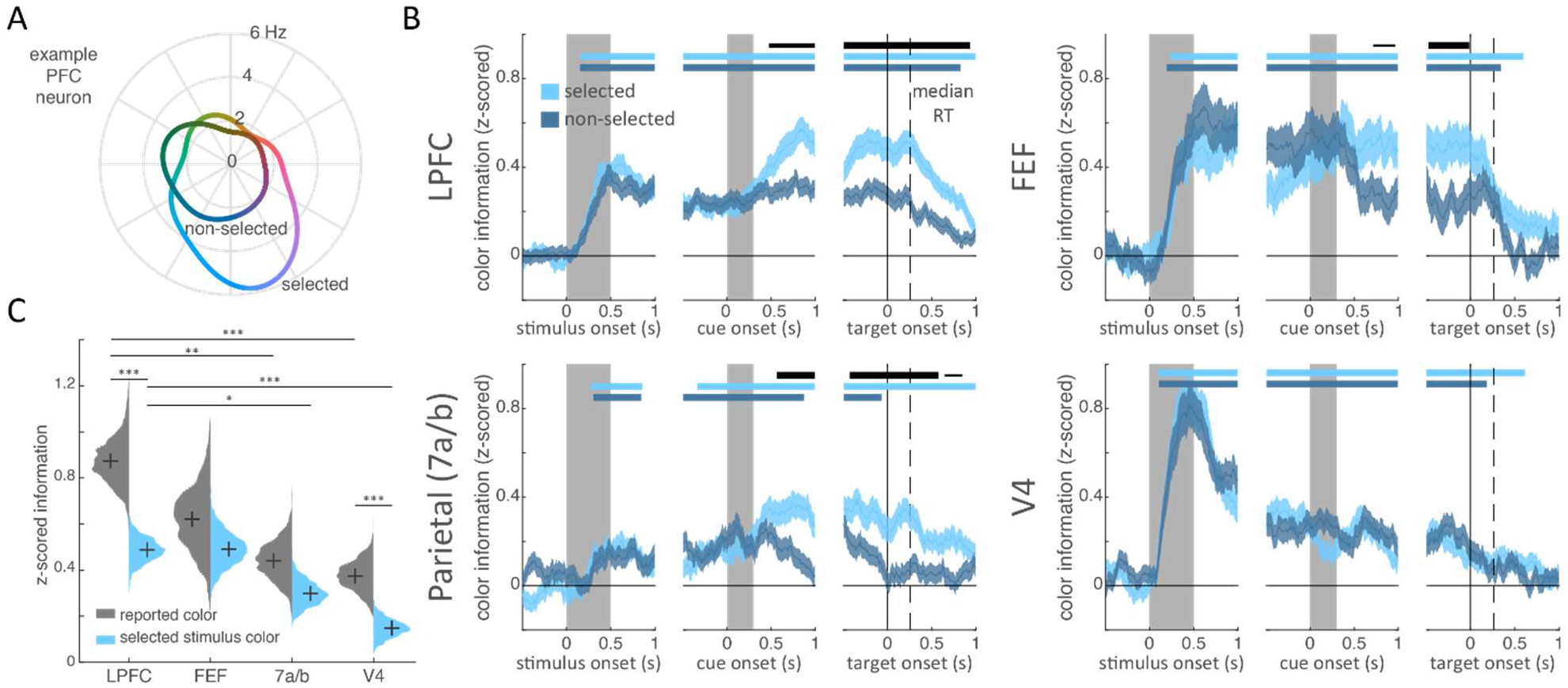
Effects of selection on color information in working memory. **(A)** Color tuning curve for an example LPFC neuron. Firing rate (radial axis) was averaged from 500-700 ms post-cue as a function of the color (angular axis) of the selected (light) and non-selected (dark) stimulus. This neuron carries information about the color of the selected item (i.e. the neural response is non-uniform), but this information is reduced for when the item is not selected. **(B)** Mean z-scored color information for the selected and non-selected color (in light and dark blue, respectively) in each brain region, averaged across all neurons. Information was quantified by calculating the entropy of the selected/non-selected tuning curves. Error bars are standard error of the mean. Horizontal bars indicate significant information for the selected item (light blue), the non-selected item (dark blue), and significant difference in information about the selected and non-selected items (black). Bar width indicate significance: p < 0.05, 0.01, and 0.001 for thin, medium, and thick, respectively. All tests were cluster-corrected for multiple comparisons (see methods). **(C)** Information about the presented color of the selected item (light blue) and the reported color of the selected item (gray), averaged across all neurons. Both types of information were calculated on firing rates in a 200 ms window prior to onset of the response color wheel. Distributions show bootstrapped estimates of the mean. Horizontal lines indicate pairwise comparisons. * p < 0.05, ** p < 0.01, *** p < 0.001.

Previous fMRI and EEG work in humans has suggested selection improves memories by enhancing the representation of a memory and/or by protecting it from decay^5,7,25^. This is thought to be similar to how attention enhances the representation of attended stimuli^31,32^; also seen here, Fig. S5). Indeed, we find selection similarly enhanced memories across prefrontal and parietal cortex. In LPFC, color information about the selected memory was increased relative to the unselected memory, starting at 475 ms after cue onset (Fig. 3B). Similar differences were seen in FEF and parietal (Fig. 3B, at 715 and 565 ms, respectively). In V4, selection did not impact memory representations across all trials (Fig. 3B) but it did have a subtle effect on trials when the animal’s memory report was more accurate (Fig. S7). In general, the selective enhancement of a memory in all four regions was related to behavior: when memories reports were inaccurate, the effect of selection was absent (in PFC and 7a) or slightly inverted (in FEF and V4), suggesting that the animal failed to select an item or selected the wrong item (Fig. S7–S8).

These results suggest selection and attention may use similar mechanisms to enhance memory/sensory representations in prefrontal and parietal cortex. However, in contrast with attention^1,31^ and motor decision-making^33^, selection did not reduce the response to the unselected memory in LPFC and parietal cortex (Fig. S9; but did slightly decrease the response in FEF), suggesting selection may not engage the competitive mechanisms that suppress unattended stimuli^32^.

Consistent with a primary role in working memory, prefrontal cortex carried more information about the color of the selected memory than both parietal and visual cortex (Fig. 3C, blue; LPFC > parietal, p = 0.029; LPFC > V4, p < 0.001; no other pairwise comparisons were significant, all by randomization test). Previous work has shown the identity of stimuli can change as they are encoded and maintained in working memory^34–38^. Reflecting this change, LPFC and V4 carried more information about the color that the animal reported at the end of the trial compared to the color of the stimulus itself (p<0.001 in LPFC and V4, randomization test). Again, relative to the other regions, prefrontal cortex carried the most information about the animal’s reported color (Fig. 3C, grey; LPFC > parietal, p < 0.001; LPFC > V4, p < 0.001; no other pairwise comparisons significant). Therefore, while we observed working memory-related signals across frontal, parietal, and visual cortices, they were strongest and most tightly linked to behavior in prefrontal cortex, particularly in LPFC.

Next, we were interested in how changing task-demands during the trial affected memory representations. Early in the trial, during the first delay, color memories must be maintained in a form that allows the animal to select the cued item (i.e. colors are bound with location information). Later in the trial, after selection, the same information is now used in a different way – to guide the visual search of the color wheel (which results in the animal’s decision and eye movement). Given this change in how memory information is used during the trial, we tested whether memory representations were transformed by selection. For these analyses, we focused on neural representations in LPFC because activity in this region encoded both stimuli and was tightly linked with behavior (Fig. 3C).

Early in the trial, before selection, the color of each item in memory was represented in separate subspaces in the LPFC neural population. Figure 4A shows the representation of color information about the upper and lower item, before selection (projected into a reduced three dimensional space, see methods for details). Color information showed a clear organization; the responses to four categories of color were separated and coded in color order for both the upper and lower item (i.e. neighboring colors on the color wheel were coded in neighboring regions of population space; note: the response wheel was rotated on each trial, so this does not reflect motor planning). Color representations for each item were largely constrained to a ‘color plane’, consistent with a twodimensional color space (these planes explained >97% of variance, Fig. 4A; see methods for details). As seen in Figure 4A, the upper and lower color planes appeared to be independent from one another, suggesting color information about the upper and lower items were separated into two different subspaces in the LPFC population (before selection).

**Figure 4.**
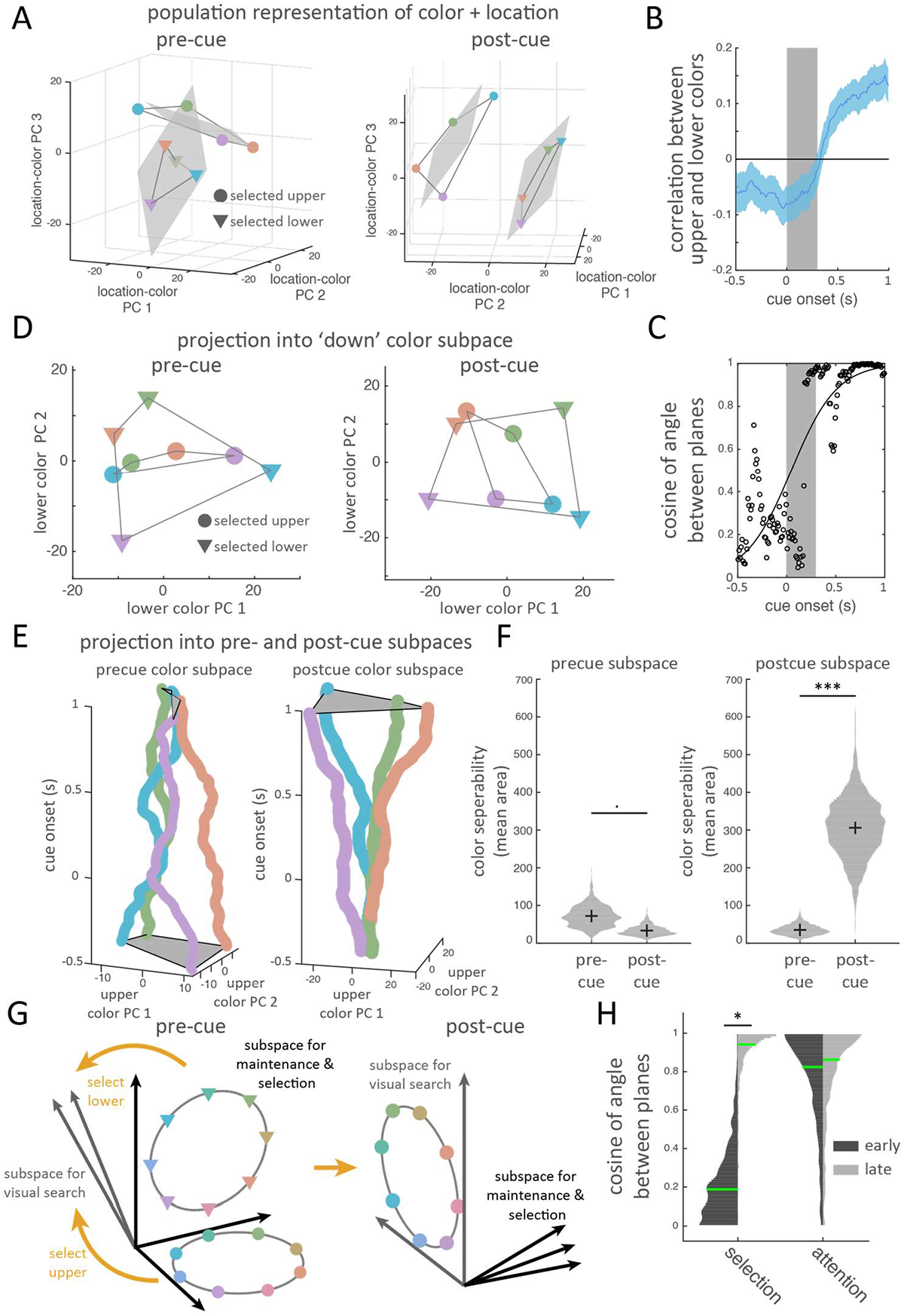
Selection transforms task-relevant information into a common subspace. **(A)** Population response for selected colors (binned into 4 color bins, indicated by marker color) at different locations (upper vs. lower, indicated by marker shape). Population response is taken as the vector of mean firing rate of all recorded neurons before the cue (pre-cue, left; taken at 400 ms) and after the cue (post-cue, right; taken just prior to target onset, see methods for details). Responses are projected into a reduced dimensionality subspace defined by the first three principle components (PCs) of all 8 color/location pairs. Grey lines connect adjacent colors along the color wheel. Gray shaded region reflects the best fitting planes to each location (see methods for details). **(B)** Color representations for upper and lower items become correlated after selection. Line shows the mean correlation between the population representation for each color when it was presented/remembered in the ‘upper’ or ‘lower’ position, over time. Correlation was measured after subtracting the mean response at each location (see methods for details). Error bars reflect standard error of the mean. **(C)** Color planes (seen in A) become aligned after selection, reflected in an increase in the cosine of the angle between the two color planes around the time of cue onset. Black line shows the best-fitting logistic function. **(D)** Alignment of color representations before (left) and after (right) selection. Colored markers indicate vector of population firing rate for both upper and lower items (markers as in A). Here, all vectors are projected into the ‘lower’ subspace, defined by the first two PCs that maximally explain variance in the color of the lower item (defined in the full N-dimensional neural space on held-out data; see methods). Timepoints and markers are as in (A). **(E)** Timecourse of population responses to the color of the upper item, projected into the upper subspace defined before selection (left) and after selection (right). Upper subspaces were defined as in D, but for the upper item. **(F)** Before selection, color representations are better separated using the pre-cue subspace. After selection, colors are better separated in the post-cue subspace. Separability was measured as the area of the quadrilateral defined by the population vectors for each color, projected into either the pre-cue or post-cue subspaces (left and right columns in each plot; area averaged across upper and lower items). Subspaces are defined as in D and E. Violin plots show bootstrapped distributions. **(G)** Schematic of how selection transforms color representations. Initially, the colors of the upper and lower item are encoded in orthogonal subspaces specific to each item’s location. The selected item is then transformed into a common subspace, regardless of its initial location. **(H)** Upper and lower representations become aligned after selection (left column) but immediately after stimulus presentation during attention (right column). Histograms show bootstrapped distribution of the cosine of the angle between the best-fitting planes for the upper and lower stimuli in either an ‘early’ (150-350 ms post-stimulus offset) or ‘late’ (200-0 ms before color wheel onset) time period during the delay. Green lines indicate median values. · p < 0.10, * p < 0.05, ** p < 0.01, *** p < 0.001.

Consistent with separate subspaces, the color representations of the upper and lower items were slightly anti-correlated with one another before selection (Fig. 4B, e.g. the N-neuron population vectors of ‘red upper’ and ‘red lower’ were anti-correlated; mean *r* = −0.067 for −300 to 0 ms pre-selection, p = 0.009, bootstrap). This weak anti-correlation suggests that tuning curves for color across the two spatial locations were slightly inverted. Further consistent with separate subspaces, the median angle between the color planes of the upper and lower items was 79.1 degrees (Fig. 4C, IQR: 71.4 to 85.1 degrees, see methods for details), suggesting these subspaces were nearly orthogonal to one another. This orthogonality was not because independent populations of neurons encoded each item: as expected from previous work^39^, more LPFC neurons were selective for both items in memory than expected by chance (31% and 35% of neurons selective for the color of upper/lower stimulus were also selective for other item; p = 1.21e-4, binomial test against proportion of doubly-selective neurons expected by chance, given the observed proportion of neurons selective to each item). Further reflecting the different upper and lower subspaces, color representations of an item were less separated when they were projected onto the *other* color subspace (e.g. projecting upper colors into the lower color subspace, Fig. 4D; each item’s color subspace was defined as the 2D space that maximally captured color information in the full N-dimensional neural space, see methods for details). To quantify the separability of colors, we measured the area of the quadrilateral defined by the four color representations (increased separation of colors increases this ‘color-area’, see methods for details). Consistent with independent subspaces for upper and lower items, color-area was greater when color representations were projected into their own color subspace compared to the other color subspace (86.1 vs. 35.2 units^2^, p=0.041, bootstrap; all subspaces were defined on held-out data).

After selection, the representation of the selected item was transformed into a different subspace (Fig. 4E). Reflecting this transformation, colors were well separated when projected into an item’s preselection subspace early in the trial, but this separation collapsed by the end of the second memory delay (Fig. 4E, left, for upper item; see Fig. S10 for lower item). Accordingly, mean color-area tended to decrease over time from 74.1 to 39.4 units^2^ (Fig. 4F, left; p = 0.076, bootstrap). In contrast, the subspace defined after selection poorly separated color responses before selection, but showed a significant increase in separation after selection (Fig. 4E, right, and S10; color-area increased from 27.8 to 261.9 units^2^ across time, Fig. 4F, right; p < 0.001, bootstrap).

Interestingly, the transformation after selection was such that the two color subspaces were now aligned (Fig. 4A). Consistent with such an alignment, the representation of the color of the selected item shifted from being anti-correlated before selection to positively correlated after selection (Fig. 4B, mean *r* = 0.1391 for −300 to 0 ms prior to target onset, p < 0.001 vs zero and vs pre-cue, bootstrap). Furthermore, the color planes of the upper and lower item shifted from orthogonal to parallel: the angle between the planes was 20.1 degrees after selection (IQR: 11.6-29.0 degrees). This change was significant over time, as measured by a significant increase in the cosine of the angle between the upper and lower color planes after selection (Fig. 4C, p = 0.006, bootstrap test of logistic regression). Finally, color representations of an item were now well separated when they were projected onto the *other* color subspace (Fig. 4D; color-area increased from 35.2 to 94.0 units^2^ over time, p=0.010, bootstrap).

The dynamic re-alignment of neural representations may reflect changing task demands during the trial. Before selection, coding of the two memories should be independent to allow for selection of a specific color (the independent subspaces of Fig. 4A, schematized in Fig. 4G, left). After selection, the color of the selected item should move into a shared ‘template’ subspace that can be used to guide visual search, regardless of whether the upper or lower item was selected (schematized in Fig. 4G, right). In this way, location information is abstracted away, as it is no longer task relevant. The alignment of upper and lower color planes may reflect this transformation (Fig. 4A,C), leading to color information about both items being represented in a similar fashion (Fig. 4B). This information can then be used to guide behavior. Indeed, from the perspective of the template space, color information is initially low (Fig. 4E, right). Then, the selection process transforms the color representation, aligning the representation of the selected color and allowing it to be read-out from the shared template space (Fig. 4A-D and 4E, right). The alignment of color spaces was important for behavior: when memory reports were inaccurate, the cosine similarity of the two color planes increased less following cue onset (Fig. S11, p = 0.0273, randomization test).

If the transformation of memories is driven by task demands, then one might expect dynamics to follow a different time course on attention trials. On attention trials, the animals can immediately prepare to search for the cued color after stimulus presentation. Accordingly, the representations of the upper and lower colors were positively correlated immediately after stimulus offset on attention trials (Fig. S12). Furthermore, the upper and lower color spaces were relatively well-aligned immediately after stimulus offset and remained aligned throughout the trial (Fig. 4H; early: median angle = 34.5°, IQR = 22.1° to 51.4°; late: median angle = 30.4°, IQR = 18.5° to 46.2°; no change with time, p = 0.449, bootstrap; there was a trend towards an interaction between time and attention/selection, p = 0.067). These results suggest the transformation of color information is under cognitive control, rather than inherent in the dynamics of the circuit.

Finally, we were interested in whether the same ‘template’ space was used in the selection and attention tasks. Consistent with a similar template space across tasks, we found a weak, but significant, correlation between color representations at the end of the delay on attention and selection trials (Fig. S13, mean rho = 0.06, p = 0.015, bootstrap). This correlation did not exist before selection (mean rho = - 0.01, p = 0.634) and increased with time (p = 0.027, bootstrap).

Altogether, our results provide novel insight into the mechanisms controlling working memory. Simultaneous recordings from prefrontal, parietal, and visual cortex show that prefrontal cortex directs selection to internal memory representations. Furthermore, selection and attention had overlapping representations in LPFC, with delayed or no generalization in FEF, parietal cortex, and visual cortex. These results suggest lateral prefrontal cortex is a ‘domain-general’ controller, categorically directing the control of representations regardless of whether they are internal memories or external stimuli. This could be useful for generalizing a task across two cognitive domains, such as sensory processing and working memory (e.g. selecting a dinner special from memory or from a printed list). Conversely, the more differentiated control signals in FEF and parietal cortex may allow for representation-specific control of memories or sensory stimuli.

Selection enhanced working memory representations in prefrontal and parietal cortex, corresponding to an improvement in working memory accuracy. This effect was similar to results seen with attention^31,32^ and when selecting a motor action^33^, although we did not observe suppression of the unselected memory. The similar changes in selectivity across behaviors suggest that there may be a common mechanism for enhancing representations, potentially to overcome interference between competing sensory inputs, motor actions, or memories.

Selection also transformed memory information. Early in the trial, working memory representations were held in location-specific ‘upper’ and ‘lower’ spaces, perhaps facilitating the selection of a memory by its associated location. Then, later in the trial, the selected memory shifted into a shared ‘template’ space, which could be used to guide responses by acting as a template for searching the color wheel. Interestingly, all three spaces (upper, lower, and template) were approximately orthogonal to one another, which could reduce interference between simultaneously maintained representations (i.e. the upper and lower stimuli) as well as limit interactions between location-bound memory representations and search-related representations.

The dynamic transformation of the memory from the upper/lower space to a shared template space is reminiscent of the rotation of motor movements from a passive ‘null’ space to an active ‘response’ space^40^. Our results build on this work, showing multiple representational spaces can converge onto a single common space (i.e. both lower and upper can transform into the template space). Furthermore, we find these dynamics are under cognitive control and depend on task demands, reflected in the different timecourses of transformation across selection and attention.

More broadly, such dynamic transformations could be a mechanism of cognitive control. Cognitive control is thought to rely on task-specific routing of information^41^. Previous work has suggested such routing can occur through gain modulation^41^ or changes in synchrony^42^-^44^. Our results suggest an alternative mechanism – cognitive control dynamically transforms information in a task-specific manner, allowing information to selectively engage with task-relevant circuits^45^. For example, consider a downstream ‘visual search’ circuit that uses color information from the common template space to guide visual search. Early in the trial, memories are stored in the upper/lower spaces, which are orthogonal to the template space (Fig. 4G, left). Thus, colors are not differentiable to the visual search circuit and so the circuit is not engaged. After selection, memory information is transformed into the shared template space (Fig. 4G, right) and the visual search circuit can be engaged. In this way, dynamically transforming representations may allow the brain to control what and when cognitive computations are engaged.

## Acknowledgements

The authors thank Britney Morea and Hannah Weinberg-Wolf for assistance with monkeys, Sina Tafazoli for assistance with microstimulation, and Flora Bouchacourt, Caroline Jahn, Alex Libby, Camden MacDowell, Sina Tafazoli, Motoaki Uchimura, and Sarah Henrickson for their feedback. We also thank the Princeton Laboratory Animal Resources staff for their support. This work was supported by NIMH R01MH115042 (TJB) and an NDSEG Fellowship (MFP).

## Materials and Methods

### Subjects

Two adult male rhesus macaques (Monkey 1 and 2, 12.1 kg and 8.9 kg) performed the experiment. All experimental procedures were approved by the Princeton University Institutional Animal Care and Use Committee and were in accordance with the policies and procedures of the National Institutes of Health.

### Behavioral task

Stimuli were presented on a Dell U2413 LCD monitor positioned at a viewing distance of 58 cm. The monitor was calibrated using an X-Rite i1Display Pro colorimeter to ensure accurate color rendering. During the experiment, subjects were asked to remember the color of either 1 or 2 square stimuli presented at two possible locations. The color of each sample was drawn from 64 evenly spaced points along an isoluminant circle in CIELAB color space. This circle was centered at (*L* = 60, a = 6, *b* = 14) and the radius was 57 units. The stimuli measured 2° of visual angle (DVA) on each side. Each stimulus could appear at one of two possible spatial locations: 45° clockwise or counterclockwise from the horizontal meridian (in the right hemifield; stimuli are depicted in the left hemifield in Figure 1 for ease of visualization) with an eccentricity of 5 DVA eccentricity from fixation. To perform the selection task, the animal had to remember which color was at each location (i.e., the ‘upper’ and ‘lower’ colors).

The animals initiated each trial by fixating a cross at the center of the screen. On selection trials, after 500 ms of fixation, one (20% of trials) or two (80% of trials) stimuli appeared on the screen. The stimuli were displayed for 500 ms, followed by a memory delay of 500 or 1,000 ms. Next, a symbolic cue was presented at fixation for 300 ms. This cue indicated which sample (upper or lower) the animal should report in order to get juice reward. Two sets of cues were used in the experiment to dissociate the meaning of the cue from its physical form. The first set (**cue set 1**) consisted of lines oriented 45° clockwise and counterclockwise from the horizontal meridian (cueing the lower and upper stimulus, respectively). The second set (**cue set 2**) consisted of a triangle or a square (cueing the lower and upper stimulus, respectively). Cues were presented at fixation and subtended 2 degrees of visual angle. After the cue, there was a second memory delay (500-700 ms), after which a response screen appeared. The response screen consisted of a ring 2° thick with an outer radius of 5°. The animals made their response by breaking fixation and saccading to the section of the color wheel corresponding to the color of the selected (cued) memory. Importantly, the color ring was randomly rotated on each trial to prevent motor planning or spatial encoding of memories. The animals received a graded juice reward that depended on the accuracy of their response. The number of drops of juice awarded for a response was determined according a circular normal (von mises) distribution centered at 0° error with a standard deviation of 22°. This distribution was scaled to have a peak amplitude of 12, and non-integer values were rounded up. When response error was greater than 60° for Monkey 1 (40° for Monkey 2), no juice was awarded and the animal experienced a short time-out of 1 to 2 s. Responses had to be made within 8 s, although, in practice, this restriction was unnecessary as response times were on the order of 200–300 ms.

Attention trials were similar to selection trials, except that the cue was presented 200-600 ms before the stimuli. After the colored squares, a single continuous delay occurred before the onset of responses screen (1300-2000 ms for Monkey 1 and 1000-2000 ms for Monkey 2). For behavioral analyses and all neural analyses around the response epoch, we only analyzed trials with a minimum of delay of 1300 ms to match the total delay range for attention and selection.

Condition (selection or attention) and cue set were manipulated in a blocked fashion. Animals transitioned among three different block types: (1) attention trials using cue set 1, (2) selection trials using cue set 1, and (3) selection trials using cue set 2. The sequence of blocks was random. Transitions between blocks occurred after the animal had performed 60 correct trials of block type 1 (attention) or 30 correct trials for block types 2 and 3 (selection), balancing the total number of attention and selection trials.

The eye position of the animals was continuously monitored at 1 kHz using an Eyelink 1000 Plus eyetracking system (SR Research). The animals had to maintain their gaze within a 2° circle around the central cross during the entire trial until the response. If they did not maintain fixation, the trial was aborted and the animal received a brief timeout.

We analyzed all completed trials, defined as any trial on which the animal successfully maintained fixation and made a saccade to the color wheel, regardless of accuracy. Monkey 1 completed 9,865 trials over 10 sessions and Monkey 2 completed 11,131 trials over 13 sessions.

As shown in Figure S1, the behavior of the two animals was qualitatively similar and so we pooled data across animals for all analyses.

### Surgical procedures and recordings

Animals were implanted with a titanium headpost to immobilize the head and with two titanium chambers for providing access to the brain. The chambers were positioned using 3D models of the brain and skull obtained from structural MRI scans. Chambers were placed to allow for electrophysiological recording from LPFC, FEF, area 7a, and V4.

Epoxy coded tungsten electrodes were used for both recording and microstimulation. Electrodes were lowered using a custom built microdrive assembly that lowered electrodes in pairs from a single screw. Recordings were acute; up to 80 electrodes were lowered through intact dura at the beginning of each recording session and allowed to settle for 2-3 hours before recording. This enabled stable isolation of single units over the session. Broadband activity (sampling frequency = 30 kHz) was recorded from each electrode. We performed 13 recording sessions in Monkey 2 and 10 sessions in Monkey 1.

After recordings were complete, we confirmed electrode locations by performing structural MRIs after lowering two electrodes in each chamber into cortex. Based on the shadow of these two electrodes, the position of the other electrodes in each chamber could be reconstructed. Electrodes were categorized as falling into LPFC, FEF, 7a, and V4 based on anatomical landmarks.

In separate experiments, we identified which electrodes were located in FEF using electrical microstimulation. Based on previous work^46^, we defined FEF sites as those for which electrical stimulation elicited a saccadic eye movement. Electrical stimulation was delivered in 200 ms trains of anodal-leading bi-phasic pulses with a width of 400 μs and an inter-pulse frequency of 330 Hz. Electrical stimulation was delivered to each electrode in the frontal well of each animal and FEF sites were identified as those sites for which electrical stimulation (<50 μA) consistently evoked a saccade with a stereotyped eye movement vector at least 50% of the time. Untested electrode sites (e.g., from recordings on days with a different offset in the spatial distribution of electrodes) were classified as belonging to FEF if they fell within 1 mm of confirmed stimulation sites and were positioned in the anterior bank of the arcuate sulcus (as confirmed via MRI).

### Signal preprocessing

Electrophysiological signals were filtered offline using a 4-pole 300 Hz high-pass Butterworth filter. For Monkey 1, to reduce common noise, the voltage time series *x* recorded from each electrode was rereferenced to the common median reference^47^ by subtracting the median voltage across all electrodes in the same recording chamber at each time point.

The spike detection threshold for all recordings was set equal to −4*σ_n_*, where *σ_n_* is an estimate of the standard deviation of the noise distribution:

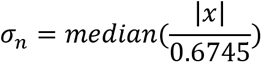

Timepoints at which *x* crossed this threshold with a negative slope were identified as putative spiking events. Repeated threshold crossings within 32 samples (1.0667 ms) were excluded. Waveforms around each putative spike time were extracted and were manually sorted into single units, multi-unit activity, or noise using Plexon Offline Sorter (Plexon, Dallas, Texas).

For all analyses, spike times of single units were converted into smoothed firing rates (sampling interval = 10 ms) by representing each spiking event as a delta function and convolving this time series with a causal half-gaussian kernel (*σ* = 200 ms).

### Mixture modeling of behavioral reports (Figures S1–S2)

Behavioral errors on delayed estimation tasks are thought to be due to at least three sources of errors^12,13^: imprecise reports of the cued stimulus, imprecise reports of the uncued stimulus, and random guessing (i.e., from ‘forgotten’ stimuli). To estimate the contribution of each of these sources of error, we used a three-component mixture model to model behavioral reports (Bays et al., 2009):

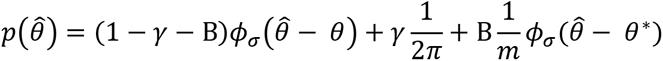

where *θ* is the color value of the cued stimulus in radians, 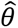 is the reported color value, *θ*\* is the color value of the uncued stimulus, *γ* is the proportion of trials on which subjects responded randomly (i.e., probability of guessing, p(Guess)), B is the proportion of trials on which subjects reported the color of the uncued stimulus (i.e., probability of ‘swapping’, p(Swap)), and *ϕ_σ_* is a von-mises distribution with a mean of zero and a standard deviation *σ* (inverse precision). Bootstrapped distributions of the maximum likelihood values of the free parameters *γ*, B, and *σ* were generated by fitting the mixture model independently to the behavioral data from each session and then resampling the best fitting parameter values with replacement across sessions. In this way, the distribution shows the uncertainty of the mean parameters across sessions.

### Calculation of cue modulation indices (Figure S3)

We developed a cue modulation index (*MI_cue_*) to describe how each neuron’s firing rate was modulated by cuing condition (‘upper’ or ‘lower’), defined as:

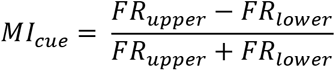

where *FR_upper_* and *FR_lower_* are a neuron’s mean firing rate on trials in which the upper or lower stimulus was cued as task relevant, respectively. Modulation indices were either computed using trials pooled across all selection trials (Fig. S3A) or calculated separately for each of the three block types (Fig. S3B-C, attention with cue set 1, selection with cue set 1, and selection with cue set 2, see above). This analysis included all neurons that were recorded for at least 10 trials per each cued location. The significance of each neuron’s modulation index (Fig. S3A) was assessed by comparing to a null distribution of values generated by randomly permuting location labels (upper or lower) across trials (1000 iterations). To test if a region had more significant neurons than expected by chance, the percentage of significant neurons was compared to that expected by chance (the alpha level, 5%).

### Classification of cued location (Figures 2 and S4)

We used linear classifiers to quantify the amount of information about the location of the cued stimulus (upper or lower) in the population of neurons recorded from each brain region (Figure 2C-D). This analysis included all neurons that were recorded during at least 60 trials for each cueing condition (upper or lower) in each block type (attention with cue set 1, selection with cue set 1, and selection with cue set 2, see above). On each of 1000 iterations, 60 trials from each cueing condition and block type were sampled from each neuron with replacement. The firing rate from those trials, locked to cue onset, was assembled into a pseudo-population by combining neurons across sessions such that pseudo-trials matched both block and cue condition. For each timestep, a logistic regression classifier (as implemented by fitclinear.m in MATLAB) with L2 regularization (*λ* = 1/60) was trained to predict the cueing condition (upper or lower) using pseudo-population data from one block (e.g., selection with cue set 1) and tested on held out data from another block (e.g., selection with cue set 2). Classification accuracy (proportion of correctly classified trials) was averaged across reciprocal tests (e.g., train on selection with cue set 2, test on selection with cue set 1).

We used a randomization test to test for significant differences in the onset time of above-chance classification across regions (Figure 2E). For each pair of regions, we computed the difference in time of first-significance (p < 0.05, bootstrap) for each region (the lag). To generate a null distribution of lags, we randomly permuted individual neurons between the two regions and then repeated the above bootstrap procedure to determine the lag in above-chance classification for each permuted dataset. 1000 random permutations were used for each pair of regions. Significance was assessed by computing the proportion of null lags of greater magnitude than the observed lag. Note that this randomization procedure controls for differences in the number of features (neurons) across regions.

To assess the discriminability of the upper and lower attention conditions (Fig. S4), we calculated the 10-fold cross validated classification accuracy (averaged across folds). To provide an estimate of variability we repeated this analysis 1,000 times, each time with a different partition of trials into folds.

### Quantification of color information (Figures 3, S5, and S9)

We adapted previous work^48^ to define a color modulation index (*MI_color_*) that describes how each neuron’s firing rate was modulated by the colors of the remembered stimuli. After dividing color space into N = 8 bins, *MI_color_* is defined as:

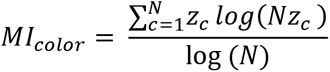

where *z_c_* is a neuron’s normalized mean firing rate *r_c_* across trials evoked by colors in the *c^th^* bin:

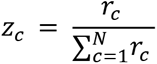

*Ml_color_* is a normalized entropy statistic that is zero if a neuron’s mean firing rate is identical across all color bins and one if a neuron only fires in response to colors from one bin. To control for differences in average firing rate and number of trials across neurons, we z-scored this metric by subtracting by the mean and dividing by the standard deviation of a null distribution of MI values. To generate this null distribution, the color bin labels were randomly shuffled across trials and the MI statistic was recomputed (1,000 times per neuron).

Z-scored color modulation indices were computed separately for each time point, trial type (attention or selection), and stimulus type (selected/non-selected/attended/non-attended, Fig. 3B and S5). This analysis included neurons that were recorded for at least 10 trials in each of these conditions. Selectivity for color was computed without respect to the spatial location of the stimulus (upper or lower). Computing selectivity for colors only presented at a neuron’s preferred location did not qualitatively change the results. Z-scored modulation indices were compared to zero or across conditions via t-test (Fig. 3B). We corrected for multiple comparisons over time using a cluster-correction^49^. Briefly, the significance of contiguous clusters of significant t-tests was computed by comparing their cluster mass (the sum of the t-values) versus that expected by chance (randomization test). Additionally, to summarize changes in selected and non-selected color information after cue onset, we averaged color information for each neuron in two time periods (−300 to 0 ms pre-cue and 200 to 500 ms post-cue) and tested the difference of these values (post-pre) against zero by bootstrapping the mean difference in color information across neurons (Figure S9).

To determine if a neuron displayed significant selectivity for the color at one particular location (upper/lower), we calculated the z-scored information about the cued color at each timepoint over the interval from 0 to 2.5 seconds post-stimulus onset independently for each location. Color selectivity was measured across all conditions, including attention, selection, and single-item trials. As described above, we used a cluster correction to correct for multiple comparisons across time. Neurons with significant color selectivity (p < 0.05) at any point during this interval were deemed color selective. Binomial tests compared the proportion of neurons with significant color selectivity for at least one of the two locations to a conservative null proportion of 10% (for two tests with an alpha of 0.05, one test for each location).

To determine if independent populations of LPFC neurons encoded the upper and lower color during the pre-cue period of selection trials, we counted the number of neurons with significant cluster-corrected selectivity during the 500 ms period before cue onset. Of the 607 neurons entering the analysis, 112 (18.5%) carried information about the upper color and 99 (16.3%) carried information about the lower color. Of these, 35 (5.8%) carried information about both the upper and lower color. Binomial tests compared this proportion (5.8%) to that expected by random assignment of top- and bottom-selectivity (i.e., 18.5% x 16.3% ≈ 3.0%).

### Quantification of reported color information (Figure 3)

To quantify the amount of information each neuron carried about the animal’s *reported* color, we followed the same approach as for stimulus color, except that responses were binned by the color reported by the animal rather than by the color of the cued or uncued stimulus (Figure 3C).

### Modulation of color information by task and behavioral performance (Figure S6–S8)

To compare the amount of color information in firing rates across the attention and selection conditions (Figure S6), we computed the z-scored color modulation indices as described above for each of the four conditions of interest (selected, non-selected, attended, and non-attended colors). Trial counts were matched across these four conditions to avoid biases in the color information statistic. To assess relative information about cued (selected and attended) and uncued (non-selected and non-attended) color information, we computed the difference in color information between each pair of conditions, for each neuron. The average difference across all neurons was then tested against zero, using the cluster correction described above to correct for multiple comparisons across time^49^.

To compare the amount of color information in firing rates when behavioral performance was relatively accurate or inaccurate (Figure S7–S8), we divided selection trials into two groups based on the accuracy of the behavioral report. Trials within each session were split by the median accuracy for that session. Z-scored color modulation indices were computed separately for each split-half of trials (Fig. S7). As above, the same number of trials were used for all four conditions (more/less accurate x selected/non-selected). Additionally, to quantify the effect of selection, the difference in color information for selected and unselected colors was computed for each group of trials separately (more or less accurate). This selected-unselected difference was then tested against zero to measure the effect of selection and tested between the two groups of trials to measure the effect of behavioral accuracy. Comparisons were done with a t-test across all neurons and used the cluster correction described above to correct for multiple comparisons across time^49^.

### Measuring the angle between upper and lower color planes (Figure 4 and S11)

As described in the main text, we were interested in understanding the geometry of mnemonic representations of color across the two possible stimulus locations (upper or lower). To explore this, we examined the population response on trials binned based on the color and location of the cued stimulus. The fidelity of these population representations depended on the behavioral performance of the animal. Therefore, for all principle component analyses, we divided trials based on the accuracy of the behavioral report (median split for each session, as above) and separately analyzed trials with lower angular error (Figure 4) and higher angular error (Figure S11).

Trials were sorted into *B* = 4 color bins and *L* = 2 locations (top or bottom), yielding *B × L = M* (8) total conditions. To visualize these population representations, we projected the population vector of mean firing rates for each of these 8 conditions into a low-dimensional coding subspace (Fig. 4A and S12C, similar to ref. 50). For each timestep, we defined a population activity matrix **X** as an *M × N* matrix, where *N* is the number of neurons:

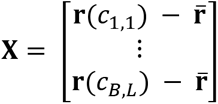

Here, **r**(*c_B,L_*) is the mean population vector (across trials) for the condition corresponding to color bin *B* and location L and 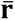 is the mean population vector across conditions (i.e., the mean of each column is zero).

The principle components of this matrix were identified by decomposing the covariance matrix **C** of **X** using singular value decomposition (as implemented by pca.m in MATLAB):

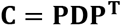

where each column of **P** is an eigenvector of **C** and **D** is a diagonal matrix of corresponding eigenvalues. We constructed a reduced (*K* = 3) dimensional space whose axes correspond to the first *K* eigenvectors of **C** (i.e., columns of **P**, **P**_*k*_, assuming eigenvectors are ordered by decreasing explained variance). These first 3 eigenvectors explained an average of 65% of the variance in the mean population response across all examined timepoints. We then projected the population vector for a given condition into this reduced dimensionality space:

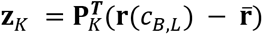

where **Z**_*k*_ is the new coordinate along axis K in the reduced dimensionality space.

We observed that, when visualized in the reduced dimensionality space, the population representations for each color bin B within a given location L tended to lie on a plane, referred to as the ‘color plane’ in the main manuscript (Fig. 4A). To identify the best fitting plane, we defined a new population activity matrix **Y**_L_ for each location L with dimensions *B* × *K*:

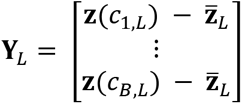

where **z**(*c_B,L_*) is the population vector for the condition corresponding to color bin B and location L in the reduced dimensionality space and 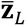 is the mean population vector across color bins for that location (i.e., the mean of each column is zero). The principle components of this matrix were calculated in the same manner as above; the first two principle components are vectors that define the plane-of-best-fit to the points defined by the rows of **Y**_L_.

If the vectors defining the plane-of-best-fit for the upper item are **v_1_** and **v_2_** and those for the lower item are **v_3_** and **v_4_**, then the cosine of the angle between these two color planes can be calculated as:

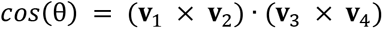

For all analyses, population vectors were based on pseudo-populations of neurons combined across sessions. Pseudo-populations were created by matching trials across sessions according to the color and location of the cued stimulus (as described above, and following ref. 39. This analysis only included neurons that were recorded for at least 10 trials for each conjunction of color and location. Confidence intervals of *cos* θ were calculated using a bootstrapping procedure. On each of 1000 iterations, 10 trials from each of the 8 conditions were sampled from each neuron with replacement. The average firing rates across these sampled trials provided the mean population vector for that condition on that iteration. To assess how *cos*(θ) changed around cue onset (Fig. 4C and S11), we used a logistic regression model of the form:

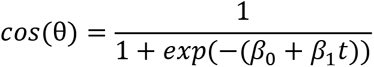

where *t* is time relative to cue onset. This model was fit to values of *cos*(θ) computed at each timepoint in the interval from 500 ms pre- to 1000 ms post-cue onset on each bootstrap iteration (described above). This yielded a bootstrapped distribution of *β*_1_ estimates which could be compared to zero or across the two groups of trials with more and less accurate behavioral responses (Figure S11).

### Defining the color subspaces for the upper and lower items in the full-dimensional space (Figure 4 and S10)

To define the color subspace in the full neuron-dimensional space, we defined *B*=*4 × N* mean population activity matrices for each location *L* in the full space:

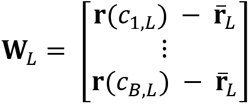

The color subspace was defined as the first two principle components of **W**_L_.

These subspaces were used for two analyses. First, we projected the population vectors of color responses from one item into the color subspace for the other item (Fig. 4D). For example, the population vector response to colors of the upper item were projected into the color subspace of the lower item, defined as the first two principal components of **W**_lower_, and vice-versa (Fig. 4D). Second, by defining the color subspace of each item at different timepoints *t_i_*, we could examine how color representations evolved during the trial (Fig. 4E-F and S10).

### Measuring separability of colors in a subspace (Figure 4 and S10)

Next, we were interested in quantifying the separability of colors in a given subspace. As seen in Figure 4D-E, the population representation of the four color conditions, projected into the subspace, form the vertices of a quadrilateral with the edges of the quadrilateral connecting adjacent colors on the color wheel (e.g., Fig. 4D). To measure separability of the colors, we computed the area of this quadrilateral (polyarea.m function in MATLAB). Bootstrapped distributions of these area estimates were obtained by resampling trials with replacement from each condition before re-computing **W**_L_.<u>Correlation of color representations (Figure 4, S12, and S13)</u>

We wanted to understand how similarly color was represented across the upper and lower locations over the course of the trial. To explore this, selection or attention trials were binned based on the color and location of the cued stimulus and then randomly partitioned into two halves. These split halves were used to estimate the degree of noise in the data (Fig S12, described below). Specifically, trials were sorted into *B* = 4 color bins, *L* = 2 locations (top or bottom), and *H* = 2 halves, yielding *B × L × H* = *M* total conditions. For each of these conditions, at a given timepoint of interest, we computed the average population vector **r**(*c_B,L,H_*).

We then computed the average correlation between each population vector and the population vectors corresponding to the same color bin at the other location (Figure 4B and S12

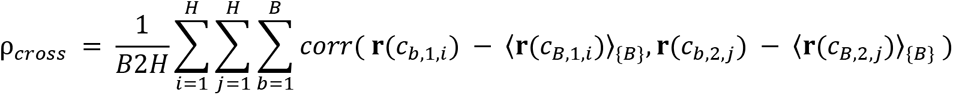

where 〈·〉_{B}_ is the average across the set of color bins *B*. In other words, for each set of B population vectors corresponding to a particular half of the data *H* and location *L*, we subtracted the mean across bins to center the vector endpoints around zero. Thus, *ρ_cross_* quantifies to what extend color representations are similarly organized around their mean across the two locations.

To obtain an upper bound on potential values of *ρ_cross_* given the degree of noise in the data, we also computed the average correlation of each population vector with itself across the two halves:

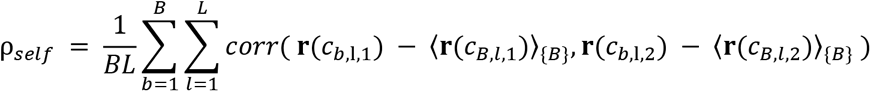

Finally, to understand how similarly color was represented across the two cueing conditions, trials were sorted into B = 4 color bins, L = 2 locations (top or bottom), and *C* = 2 cuing conditions (attention/selection). For each of these conditions, at a given timepoint of interest, we computed the average population vector **r**(*c_B,L,C_*). We then computed the average correlation between each population vector and the population vectors corresponding to the same color bin at either the same or different location in the other task (Fig. S13):

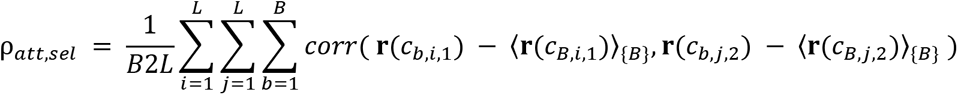

To compare the similarity of color representations on selection trials to pre-target attention color representations, we computed this correlation between (1) the response on attention trials, for all timepoints falling within the interval from −300 ms to 0 ms before the onset of the response wheel, and (2) the response on selection trials at two different timepoints: before selection (from −300 to 0 ms before the cue) and after selection (from −300 to 0 ms before the onset of the response wheel). Correlation was measured between each timepoint across windows and then averaged across all pairs of timepoints.

As above, population vectors were pseudo-populations of neurons combined across sessions, where trials across sessions were matched according to color bin and location^39^. This analysis only included neurons that were recorded for at least 10 trials for each conjunction of color and location. Confidence intervals for *ρ_cross_*, *ρ_self_*, and *ρ_att,sel_* were calculated with a bootstrap. On each of 1000 iterations, and for each neuron and condition (color–location–half conjunction), the entire population of trials in that condition was resampled with replacement. The average firing rates across these sampled trials provided the mean population vector for that condition on that iteration. As with principle components analyses, we divided trials based on the accuracy of the behavioral report (median split of trials for each session) and the presented results reflect analysis of trials with lower angular error, unless otherwise noted.

## Supplementary Figures

**Figure S1.**
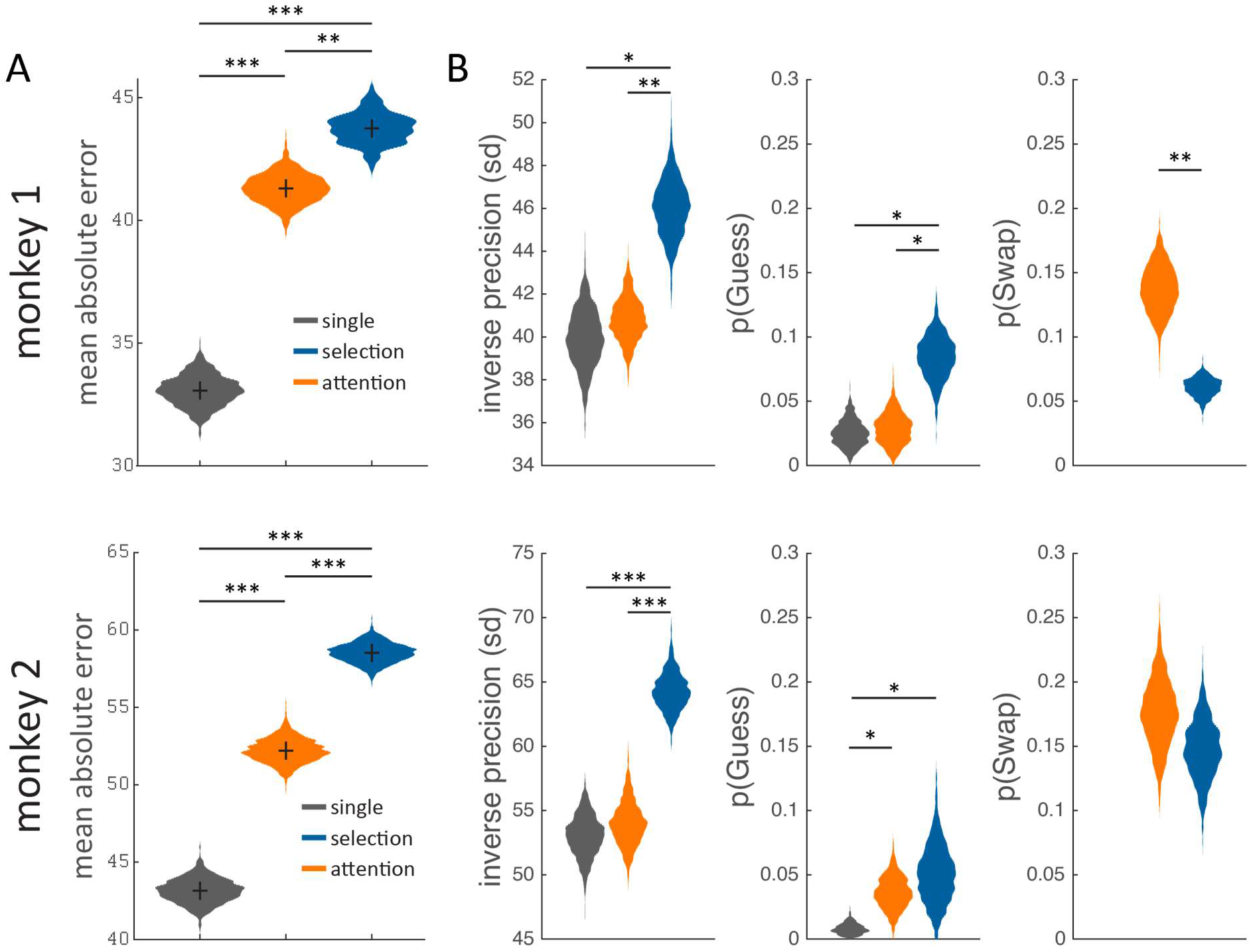
**(A)**Mean absolute angular error and **(B)** mean mixture model parameter fits for each animal. Violin plots depict bootstrapped distribution. * p < 0.05, ** p < 0.01, *** p < 0.001. Although monkey 1 displayed slightly better performance than monkey 2, the animals displayed similar patterns of performance across conditions.

**Figure S2.**
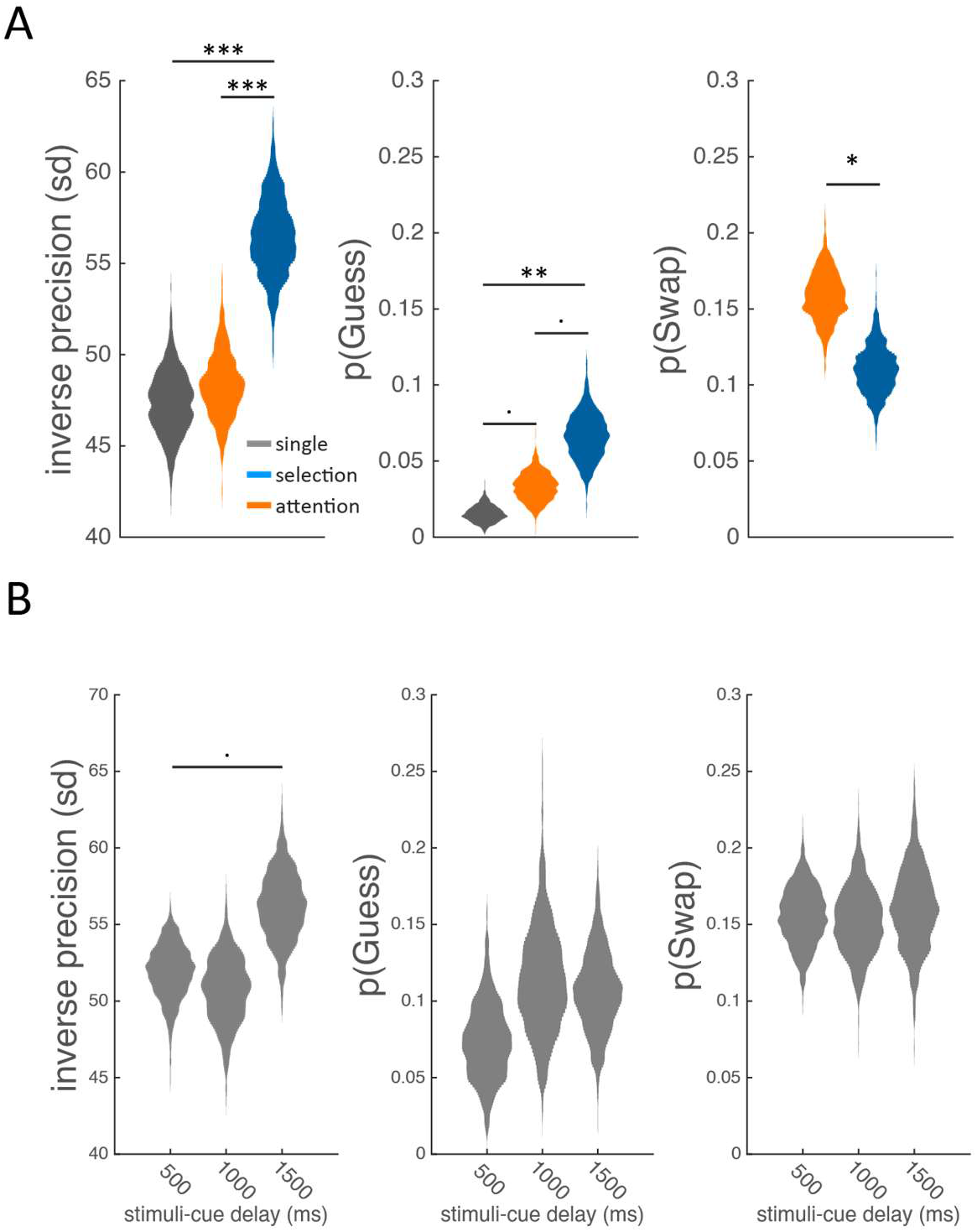
Mixture model parameter fits of behavior pooled across animals for the **(A)** main task shown in Figure 1A and **(B)** the retro-cue timing manipulation shown in Figure 1D. · p < 0.1, * p < 0.05, ** p < 0.01, *** p < 0.001.

**Figure S3.**
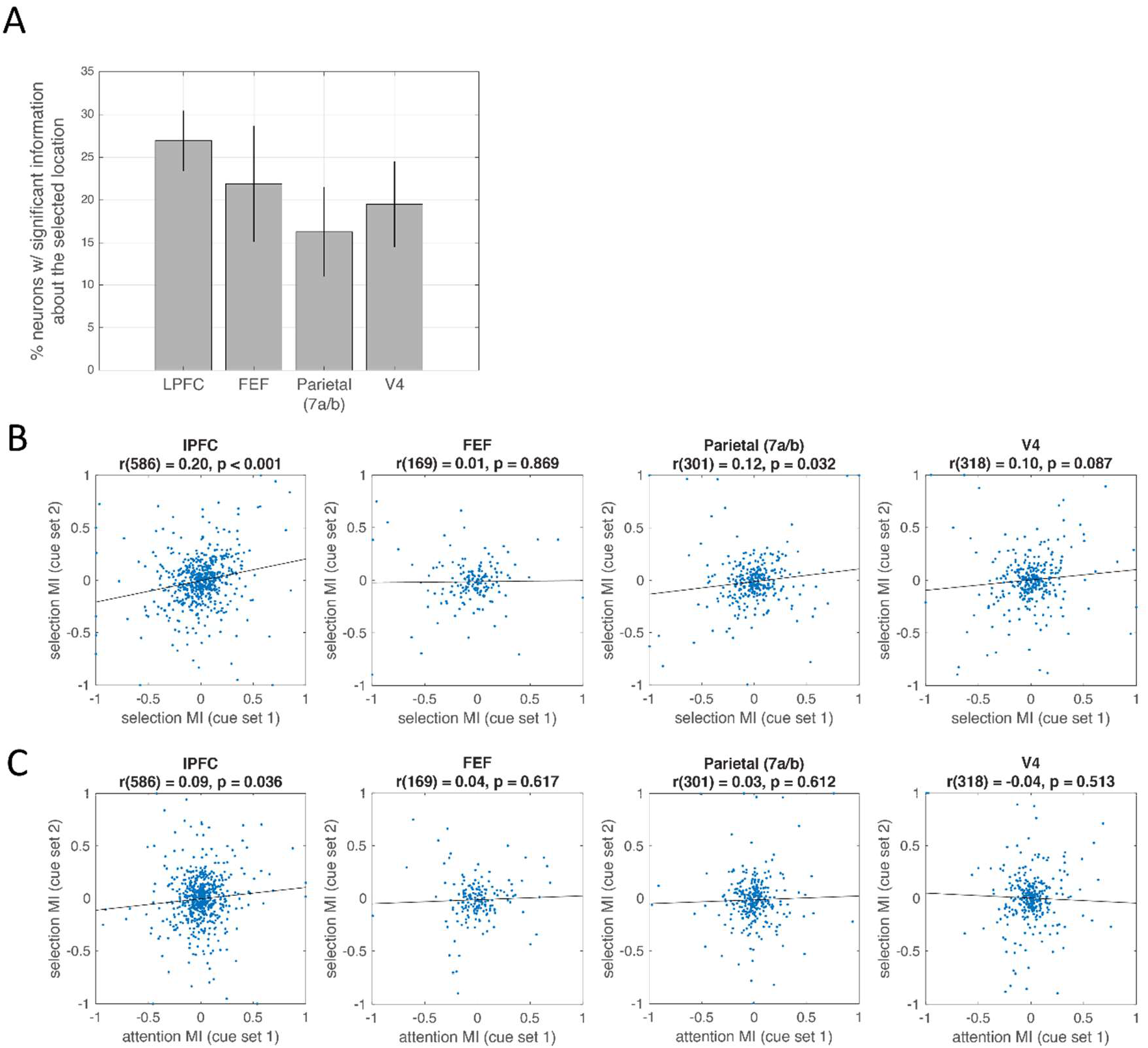
**(A)** Percent of neurons in each region of interest. with firing rates that were significantly modulated by the selected location after cue onset (trials pooled across cue set 1 and 2). For each neuron, we quantified location selectivity using a modulation index (see methods) and compared this value to a null distribution by permuting location labels across trials. All four regions showed strong selectivity: LPFC had 159 out of 590 neurons selective; FEF: 37/169, 7a/b: 49/301, V4: 62/318, all p < 0.001 for binomial test against chance of 5%. **(B)** Correlation of modulation indices for selection cue set 1 and 2. Positive modulation indices indicate that a neuron displayed a higher firing rate in response to the “upper” cue within a particular cue set, and negative modulation indices indicate that a neuron displayed a higher firing rate in response to the “lower” cue within a particular cue set. Positive correlations indicate that neurons coded for the selected location in a consistent fashion across cue sets. **(C)** Correlation of modulation indices for selection cue set 2 and attention cue set 1. Positive correlations indicate that neurons coded for the selected location in a consistent fashion across cue sets and conditions (attention/selection).

**Figure S4.**
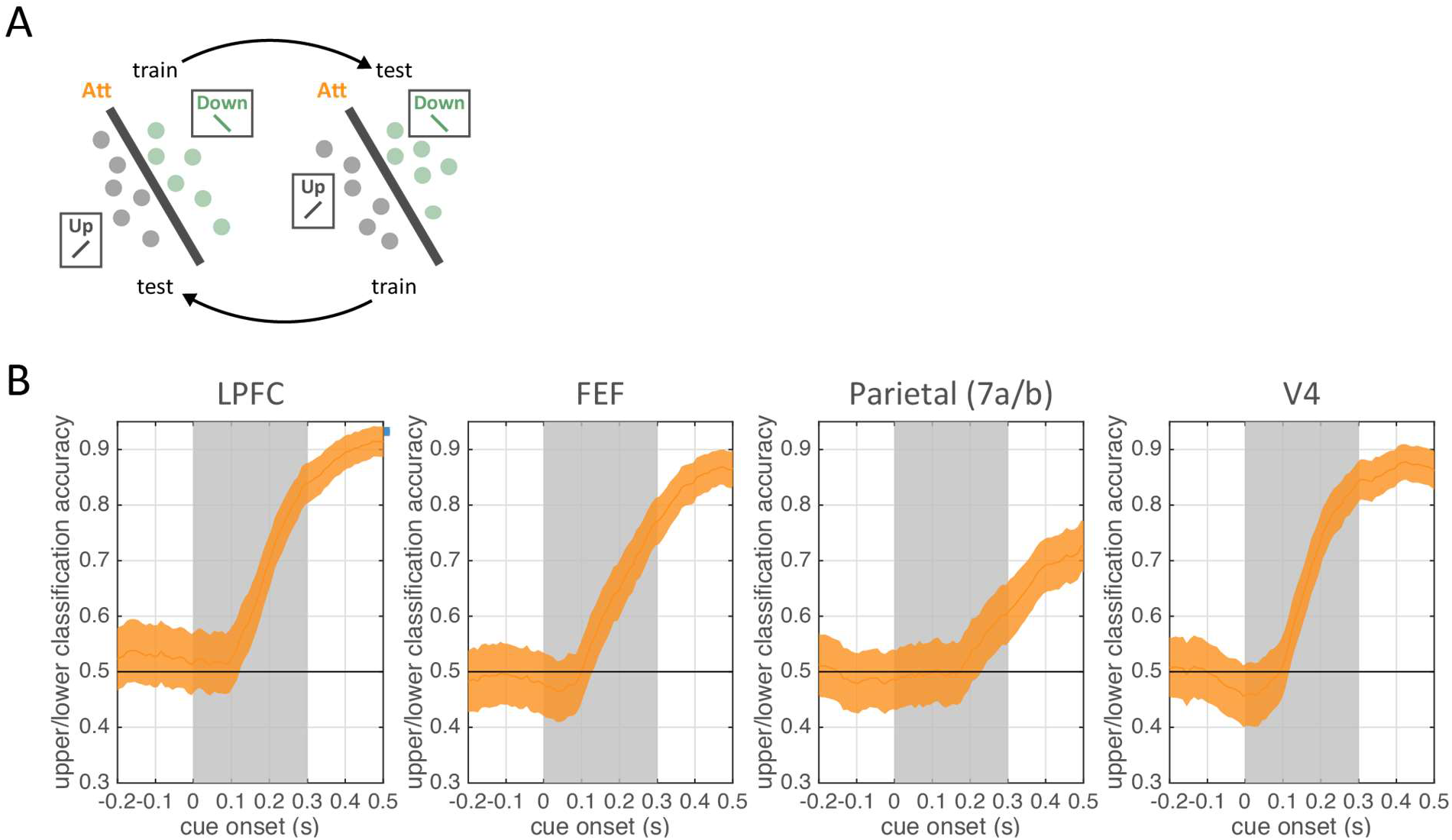
**(A)** To confirm that the neural responses to the two cue conditions were discriminable on attention trials, we calculated the cross-validated classification accuracy (10-fold, one example fold shown for clarity). **(B)** Mean classification accuracy for each brain region, relative to cue onset. Error bars reflect the standard deviation across 1,000 iterations, each with a different partition of trials into folds. Note that this analysis does not reveal the timecourse of attentional control because it confounds information about cued location (up or down) and the visual appearance of the cue itself (a line oriented +/- 45 degrees). Rather, these results indicate that these two conditions are separable in all brain regions, and so failures in cross-classification performance (Fig. 2D, purple traces) cannot be ascribed to poor separability of these two conditions.

**Figure S5.**
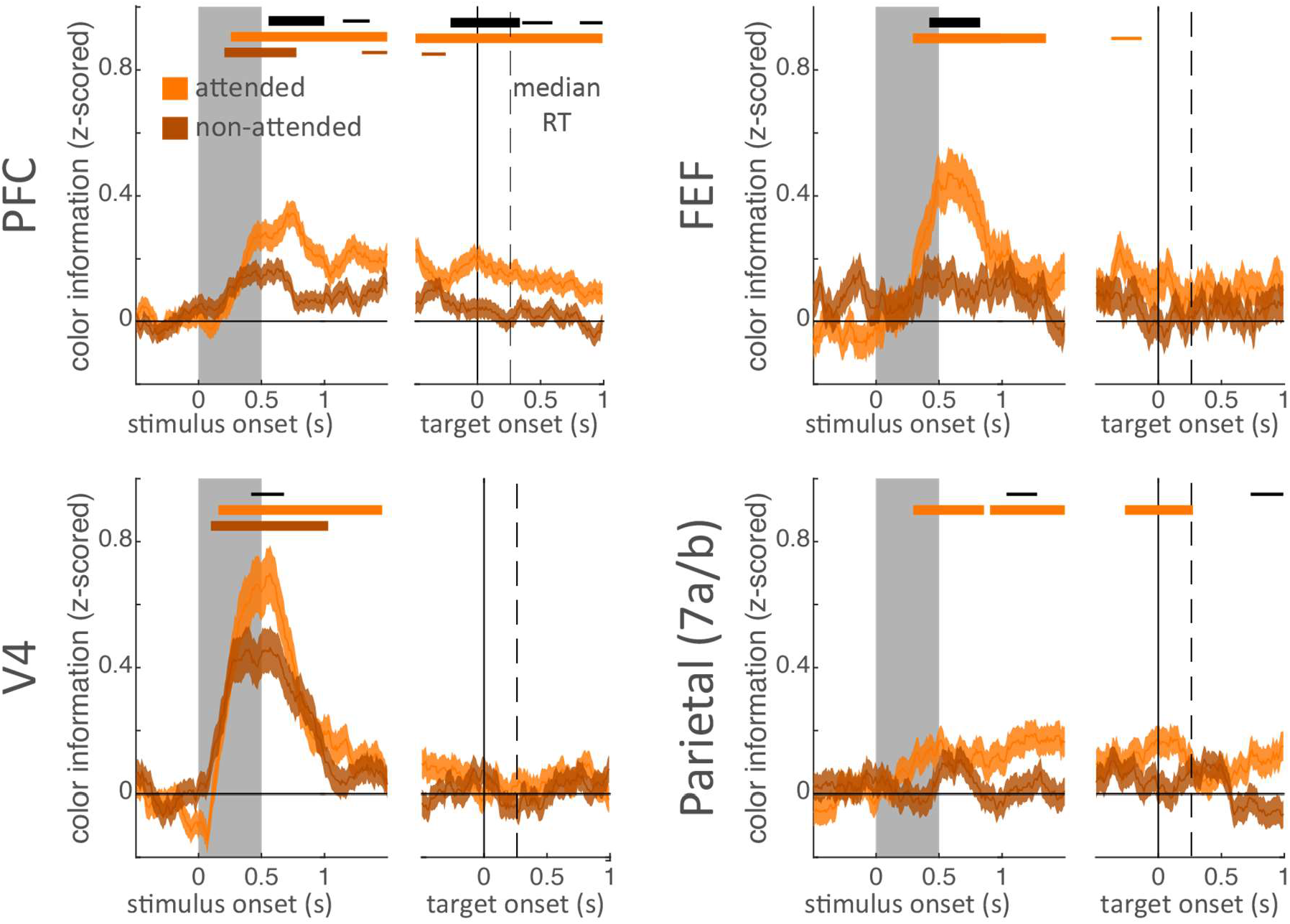
Mean z-scored color information for the attended and non-attended color on attention trials. Error bars are standard error of the mean. Horizontal bars indicate significant information for the attended item (light orange), the non-attended item (dark orange), and significant differences in information about the attended and non-attended items (black). Bar width indicate significance: p < 0.05, 0.01, and 0.001 for thin, medium, and thick, respectively.

**Figure S6.**
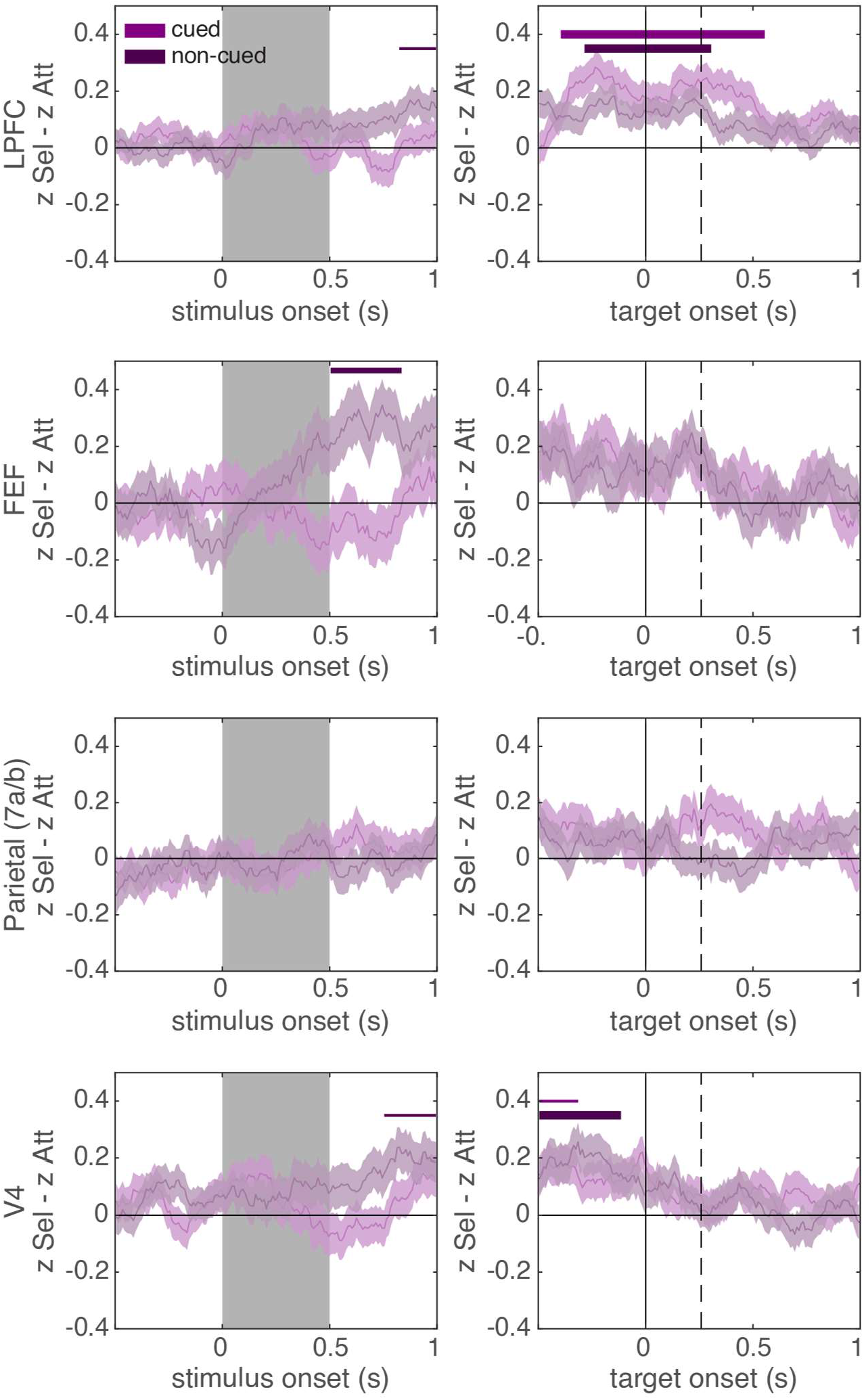
Difference in z-scored color information between selection and attention trials for the cued item (selected/attended; light purple) and uncued item (non-selected/non-attended; dark purple). Positive values indicate there was more information about an item on selection trials. Error bars are standard error of the mean, created by bootstrapping across cells (see methods for details). Horizontal bars indicate significant differences from zero (i.e. differences between selection and attention) for the cued item (light purple) and the non-cued item (dark purple). Bar width indicate significance: p < 0.05, 0.01, and 0.001 for thin, medium, and thick, respectively.

**Figure S7.**
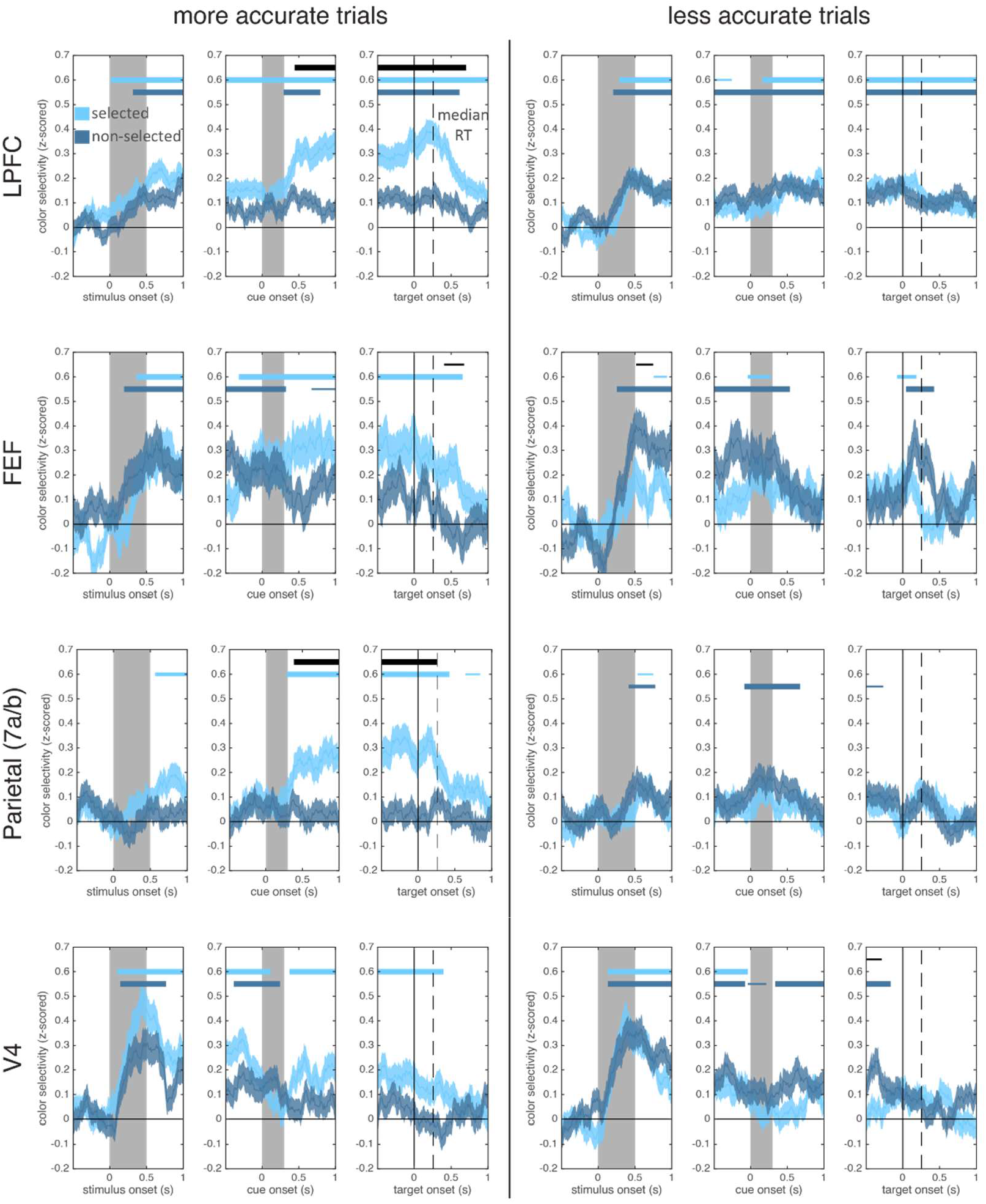
Mean z-scored color information for the selected (light blue) and non-selected color (dark blue), separated into trials with more accurate behavioral responses (left column; error was less than median error) and less accurate behavioral responses (right column; error was greater than median error). Plots followed Figure 3B. Horizontal bars indicate significant information for the selected item (light blue), the non-selected item (dark blue), and significant differences in information about the selected and non-selected items (black). Bar width indicate significance: p < 0.05, 0.01, and 0.001 for thin, medium, and thick, respectively.

**Figure S8.**
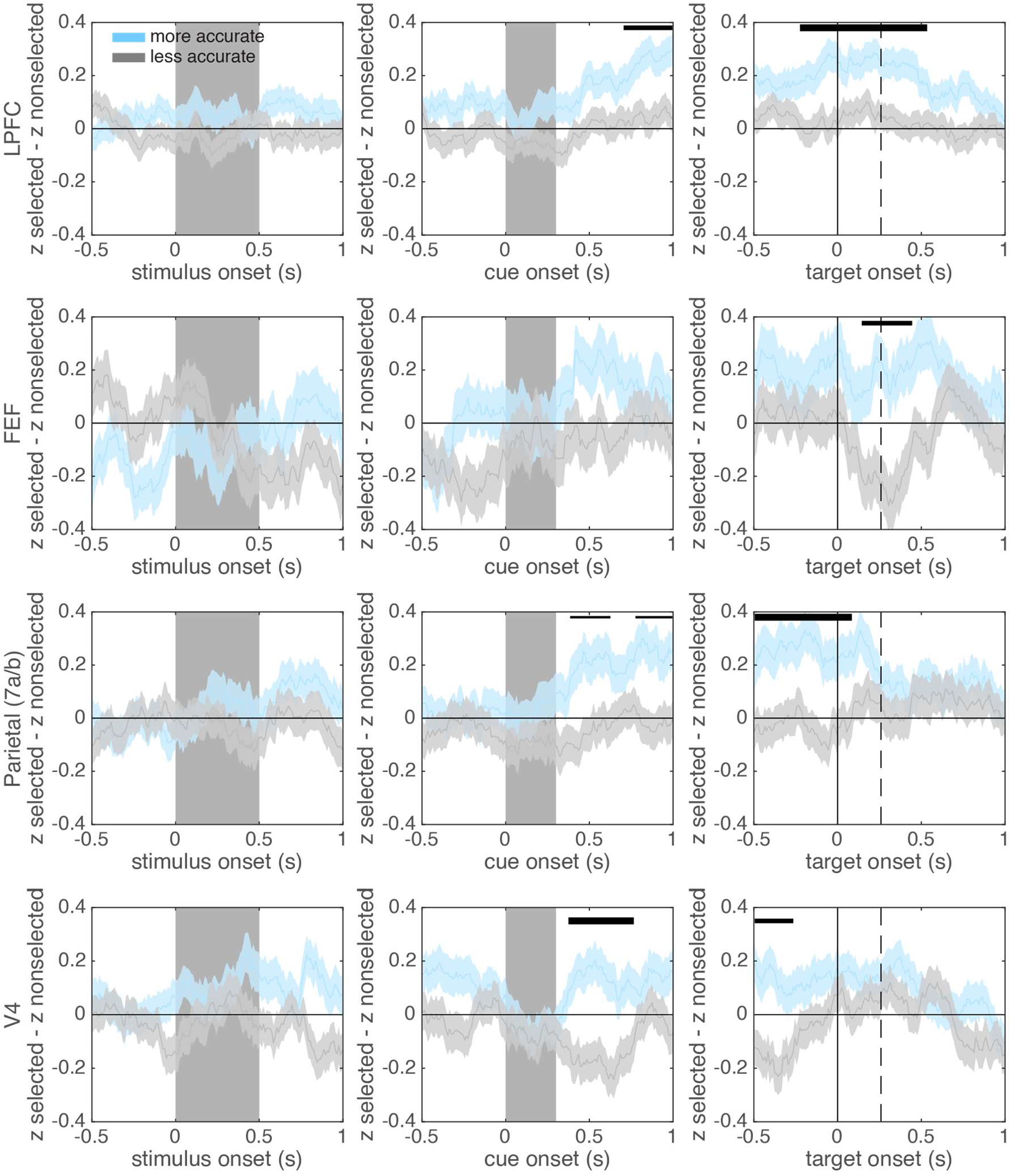
Difference in z-scored color information between the selected and non-selected item for more accurate and less accurate trials. As in Figure S7, trials were split based on angular error (relative to median error). Positive values indicate there was more information about the selected item than the non-selected item. Error bars indicate standard error of the mean. Horizontal bars indicate significant differences between more and less accurate trials. Bar width indicate significance: p < 0.05, 0.01, and 0.001 for thin, medium, and thick, respectively.

**Figure S9.**
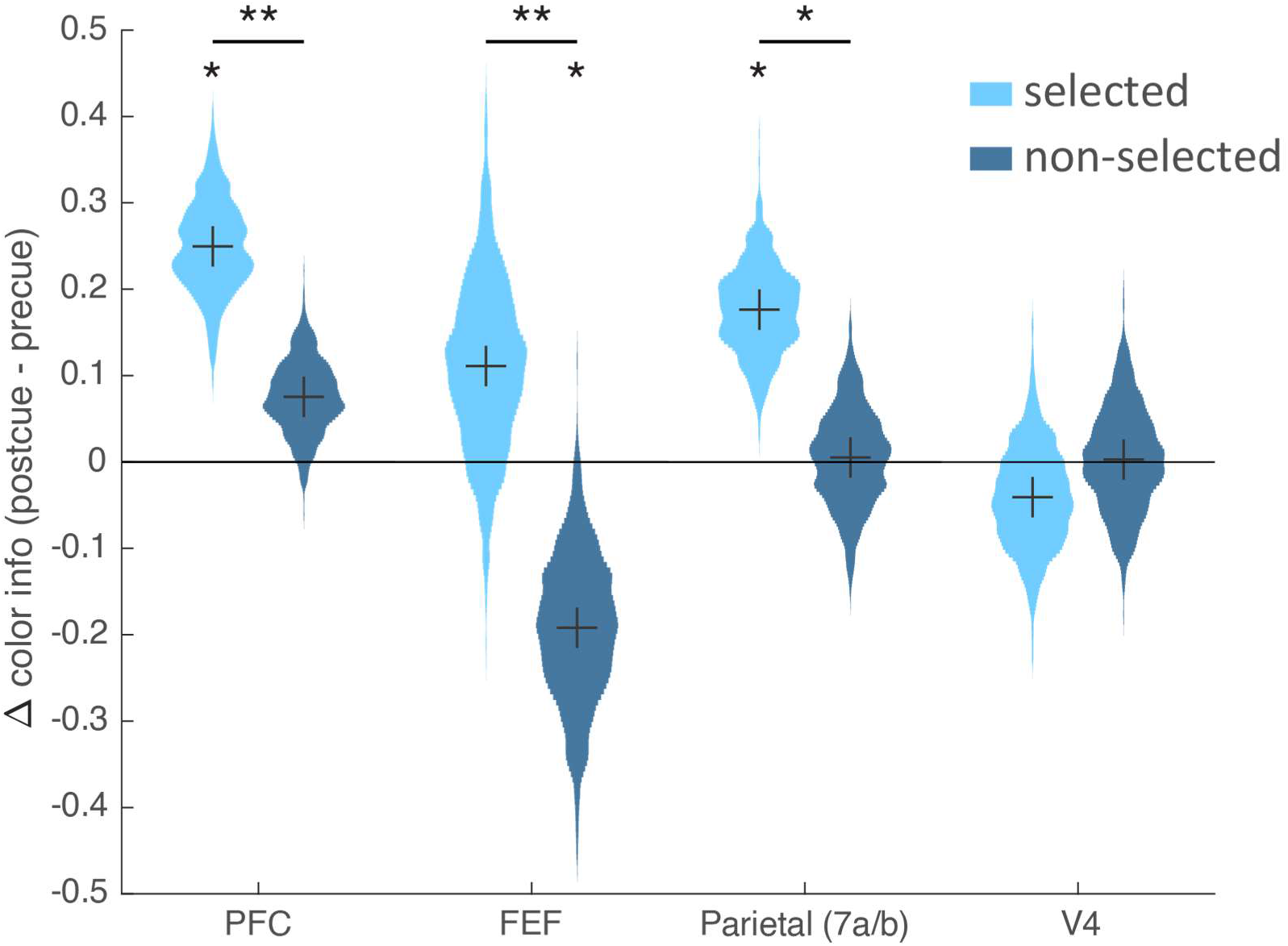
Selection enhanced the representation of the selected item in frontal and parietal regions and reduced the representation of the un-selected item in FEF. Y-axis shows the increase in color information after selection (post-cue period: 200 to 500 ms after cue offset), relative to information before selection (pre-cue period: −300 to 0 ms before cue onset). Violin plots show the distribution of this difference (bootstrapped across neurons). * p < 0.05, ** p < 0.01, *** p < 0.001.

**Figure S10.**
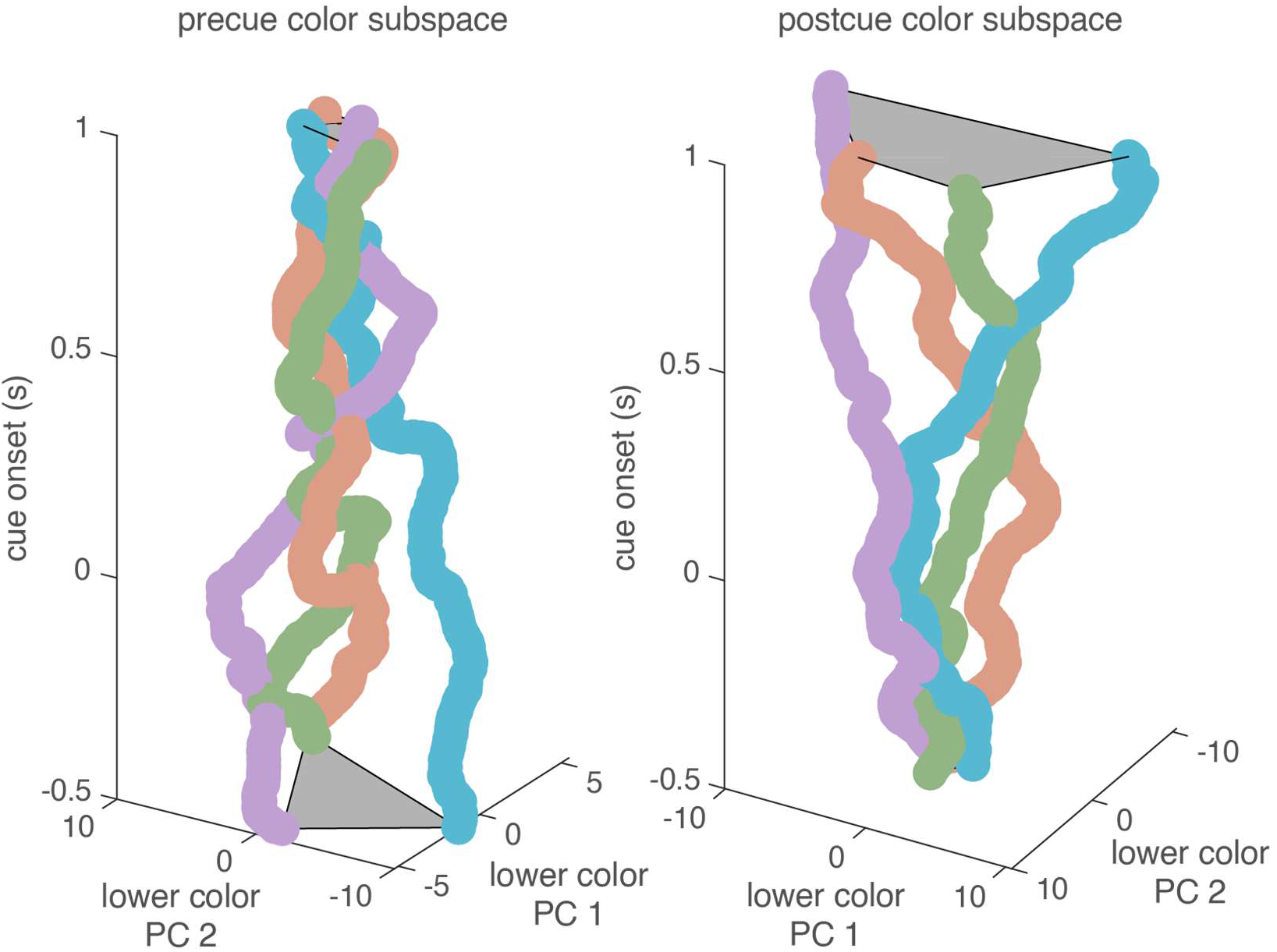
Population trajectories for ‘lower’ colors, over time, as projected into the ‘lower’ color subspace. Lower color subspace was defined as a 2D space that maximally explained variance across the four ‘lower’ colors (see methods for details). Subspace was defined either before or after selection, as in Figure 4E. As for the ‘upper’ color (Fig. 4E), temporal cross generalization was poor, suggesting the color information was represented in a different subspace by the end of the trial.

**Figure S11.**
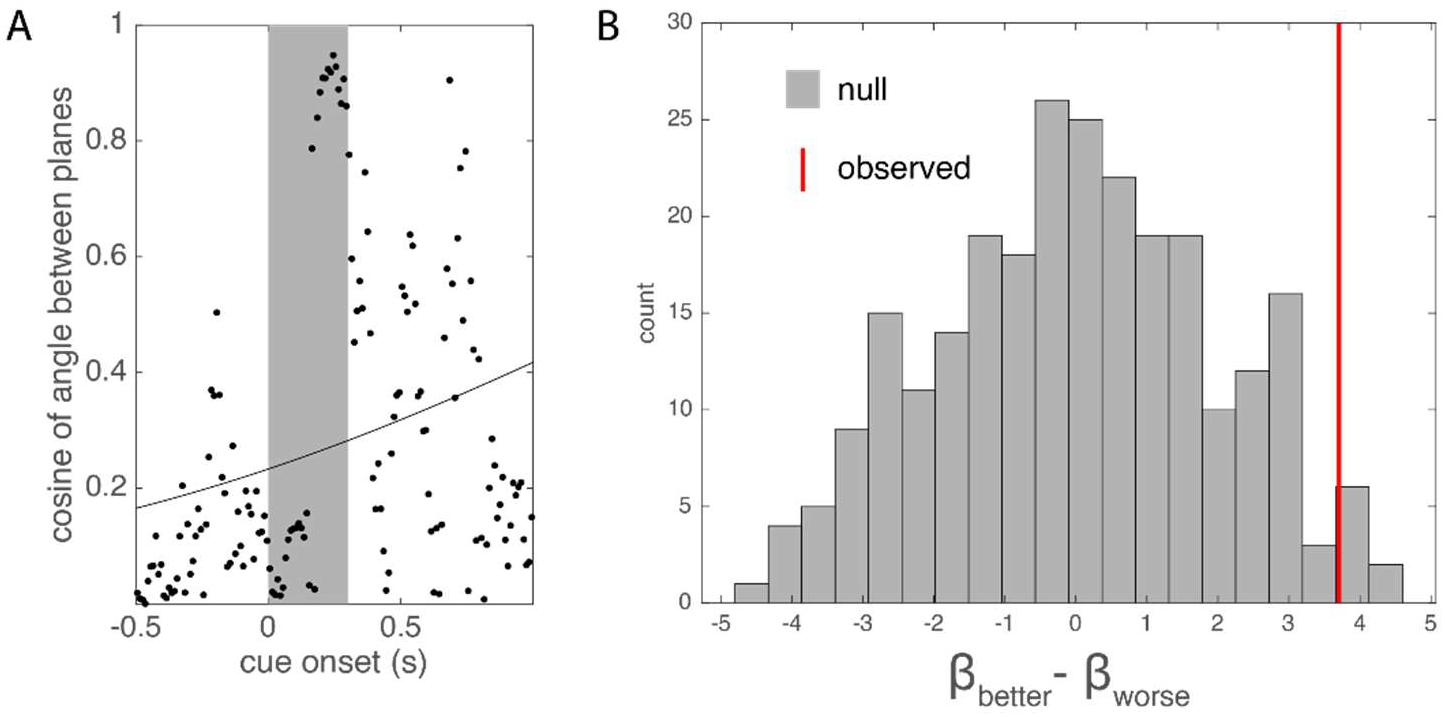
**(A)** The upper and lower color planes do not align on inaccurate trials. Figure follows Figure 4C, but shows data for trials in which absolute angular error was greater than the median error. Black markers show the cosine of the angle between the two color planes around the time of cue onset. Black line shows best-fitting logistic function. **(B)** Difference in the slope of the logistic fit (i.e., the coefficient for time, see methods) between trials in which angular error was ‘better’ or ‘worse’ than the median (red line). Null distribution (gray histogram) was generated by randomly permuting the accuracy label (‘better’ or ‘worse’) between population vectors for the same color-location conjunction. Trials in which the animal was more accurate were associated with a greater increase in the cosine of the plane angle (i.e. a greater increase in alignment) around cue onset.

**Figure S12.**
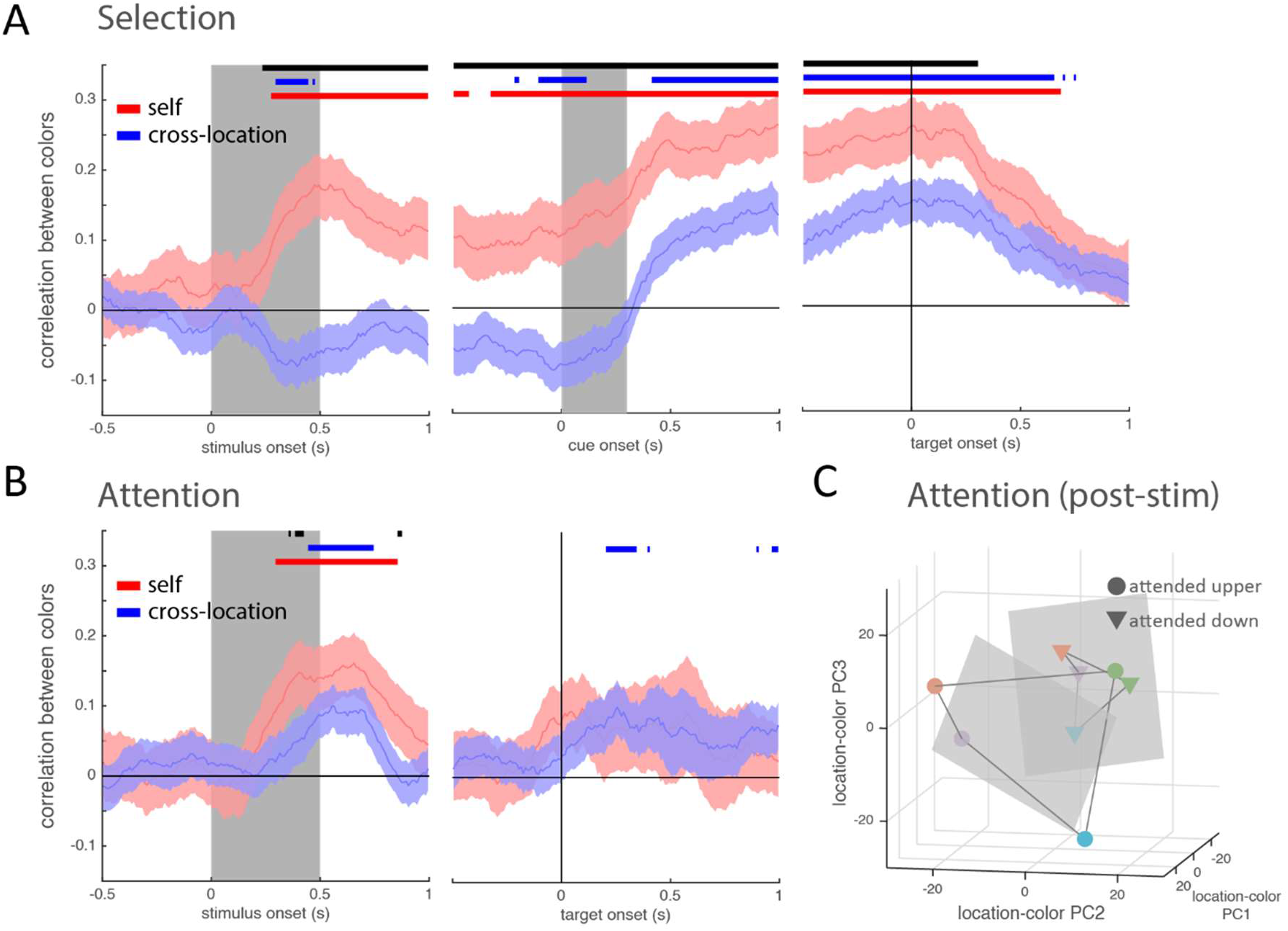
**(A)** Correlation of population vectors representing colors at the same location (self; red line) or between locations (cross-location; blue line) on selection trials. Correlations were measured after subtracting the mean vector at each location (as in Figure 4B; see methods for details). Self-correlation was computed on held-out data and provides an upper-bound on the between-location correlation values, given the noise level. Bars reflect uncorrected t-tests (p < 0.05) for each correlation type vs zero (red and blue) and between each other (black). **(B)** Same as in A, but for attention trials. **(C)** Population responses 200 ms after stimulus offset on attention trials (projected into a reduced dimensional space for visualization). Markers indicate mean position of population activity for each condition (binned by the color and location of the attended item) in a subspace spanned by the first three principle components that explain the most variance across all 8 conditions.

**Figure S13.**
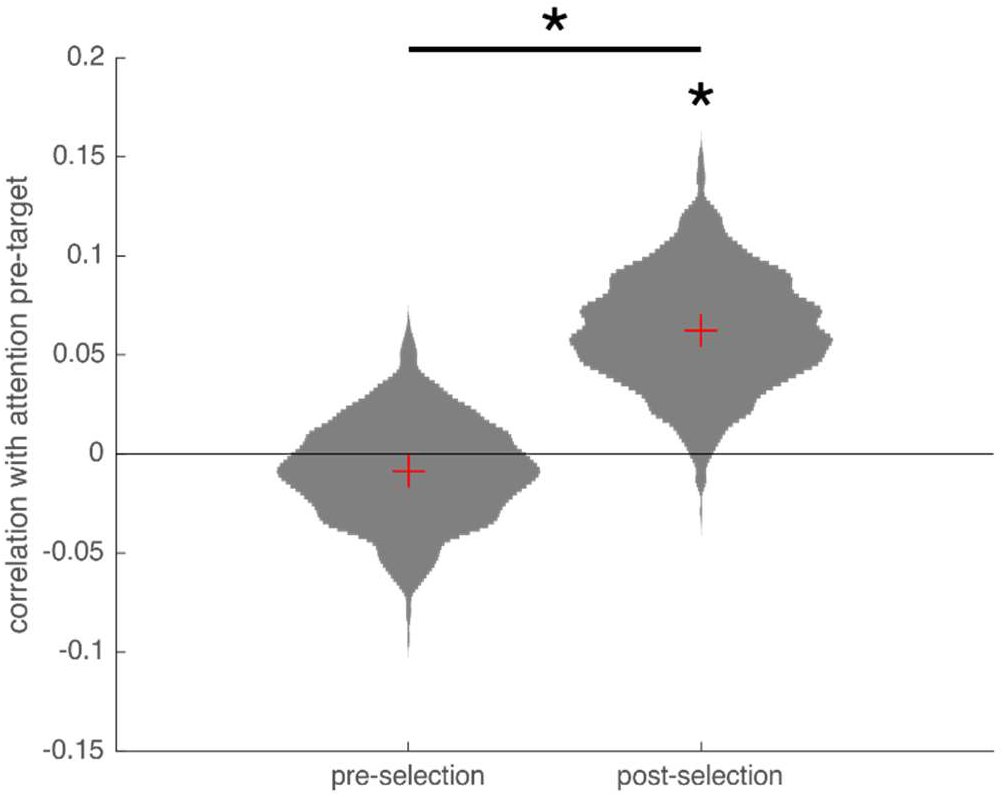
Mean correlation between the population representations for each color across attention and selection tasks (after subtracting the mean vector at each location, as in Fig. 4B and S12, see methods for details). Violin plots reflect the bootstrapped estimate of the distribution about the mean. The mean correlation was computed between the color representations taken from the 300 ms before the onset of the response wheel on attention trials and the color representations taken from either a pre-selection period (left distribution; −300 to 0 ms pre-cue) or post-selection period (right distribution; - 300 to 0 ms before response wheel onset).

